# Characterization of the Kaposi’s sarcoma-associated herpesvirus terminase complex component ORF29

**DOI:** 10.1101/2024.11.20.624617

**Authors:** Yuki Iwaisako, Youichi Suzuki, Takashi Nakano, Masahiro Fujimuro

**Affiliations:** Department of Cell Biology, Kyoto Pharmaceutical University, Misasagi-Shichono-cho 1, Yamashina-ku, Kyoto, 607-8412, Japan; Department of Microbiology and Infection Control, Faculty of Medicine, Osaka Medical and Pharmaceutical University, Osaka, 569-8686, Japan

## Abstract

Kaposi’s sarcoma-associated herpesvirus (KSHV) belongs to the gammaherpesvirinae subfamily. In the lytic phase of herpesviruses, viral capsids are formed in the nucleus of the host cell and the replicated viral genome is packaged into capsids. The herpesviral genome is replicated as a precursor head-to-tail concatemer consisting of tandemly repeated genomic units, and each genomic unit is flanked by terminal repeats (TRs). The herpesvirus terminase complex packages one genomic unit into a capsid, by cleavage of the TRs in a precursor genome. Although the terminase complexes of alpha- and beta-herpesviruses are well characterized, in KSHV, the terminase complex is poorly understood. KSHV ORF7, ORF67.5, and ORF29 are thought to be components of the terminase complex. We previously reported that ORF7- or ORF67.5-deficient KSHV formed immature soccer ball-like capsids and failed to cleave the TRs, resulting in decreased virion production. Moreover, ORF7 interacted with ORF29 and ORF67.5. However, ORF29 and ORF67.5 did not interact with each other. Thus, although ORF7 and ORF67.5 are important for KSHV terminase function, the function of ORF29 remains largely unknown. Here, we constructed ORF29-deficient BAC16 and analyzed its virological properties. ORF29 was essential for virion production and TR cleavage. Numerous immature soccer ball-like capsids were observed in ORF29-deficient KSHV-harboring cells. The N-terminal region of ORF29 was important for its interaction with ORF7, although full-length ORF29 was required for effective complexation of the KSHV terminase. Furthermore, ORF29 preferentially interacted with itself rather than with ORF7. Thus, our data shows that ORF29 functions as a fundamental terminase component.

**IMPORTANCE:** Since the role of ORF29 in the KSHV terminase complex remains unknown, we constructed ORF29-deficient KSHV. Our results demonstrated that ORF29 functions as a component of the KSHV terminase and is critical for capsid formation, TR cleavage, and terminase complexation. Moreover, ORF29 robustly interacted with itself. In HSV-1, a terminase complex containing UL15, UL28, and UL33 forms a trimer, and six trimers assemble into a hexameric ring. The HSV-1 genome passes through this ring and undergoes TR cleavage and genome packaging into a capsid. The self-interaction of ORF29 may be involved in the multimerization of a terminase complex or formation of the KSHV terminase ring. In addition, a novel KSHV protein, which we refer to as ORF29.5, was expressed during the lytic state. ORF29.5 mRNA shares the same stop codon with the ORF29 mRNA. Moreover, a part of the ORF29.5 coding region may overlap with the C-terminal ORF29 coding region.

## Introduction

KSHV, also known as human herpesvirus 8 (HHV-8), causes Kaposi’s sarcoma and primary effusion lymphoma in AIDS patients (1). When KSHV infects healthy individuals, it maintains a latent infection in vascular endothelial cells or B cells. In the latent infection state, KSHV latent genes are expressed, and these latent gene products contribute to cell proliferation, anti-apoptosis, stabilization of the viral genome, and maintenance of KSHV latency. In KSHV-latently infected individuals, reactivation of KSHV is induced by UV exposure, immunodeficiency, drug administration, or hormonal changes. Reactivation of KSHV induces the transition to lytic infection, which is characterized by progeny virus production (2). In the KSHV lytic infection cycle, lytic gene expression, viral genome replication, capsid formation, viral particle formation, and budding occur sequentially. KSHV lytic genes are classified as immediate-early (IE), delayed-early (DE), and late (L) genes based on their expression timing and expression requirements (2). The mechanisms of KSHV precursor genome synthesis, precursor genome processing, genome insertion into the capsid, capsid maturation, and infectious virus particle (i.e., virion) formation are largely unknown. However, these mechanisms have been hypothesized to include the following steps, which are described below. Initially, the KSHV DNA genome is replicated as head-to-tail concatemers. A head-to-tail concatemer consists of a linear form of tandemly repeated genomic units, and each genomic unit is flanked by terminal repeat sequences (TRs) (3). A single TR comprises a G/C-rich sequence of 801 base pairs (bps) and a set of TRs is composed of 20-40 tandemly linked TRs. The one unit-length viral genome is flanked by a set of 20-40 tandemly linked TRs. It is speculated that the viral genome is packaged into a capsid, and the one unit-length viral genome is generated by cleavage at the TR portion within the viral genome precursor. The order of genome packaging and genome cleavage is unclear, but these processes are mediated by the viral terminase complex. However, little is known about the processing of the KSHV precursor genome and the insertion of the processed genome into a capsid by the terminase complex. These processes result in the formation of a mature capsid. The mature capsids acquire viral tegument proteins and lipid bilayer envelopes and egress from the host cell as virions.

Compared to the terminase complex of KSHV, that of herpes simplex virus 1 (HSV-1) is better understood. The HSV-1 terminase complex is responsible for packaging of the viral genome into the capsid lumen, cleaving the TR portion within the viral genome precursor, and producing one unit-length viral genomes. Furthermore, the HSV-1 terminase complex has been shown to have motor activity that moves the DNA strand in an ATP-dependent manner to facilitate viral genome packaging into the capsid. The HSV-1 terminase complex also recognizes the specific DNA sequence for TR cleavage (4). On the other hand, only a few papers have been published on the terminase complex of KSHV. Based on homology with other herpesviruses, ORF7, ORF29, and ORF67.5 of KSHV are considered to be components of the terminase complex (5). We have previously reported that ORF7 and ORF67.5 are required for virus production and are essential for TR cleavage (6, 7, 8). KSHV lacking ORF7 or ORF67.5 failed to form mature capsids and instead formed soccer ball-like capsids, which are thought to be immature capsids (6, 7, 8). However, the function of ORF29 as a component of the KSHV terminase complex remains unknown. Therefore, in this paper, we focused on ORF29 and characterized its role as part of the KSHV terminase complex.

The terminase functions of KSHV ORF29 remain largely unclear, although the HSV-1 homolog of ORF29 (UL15) is well understood. The coding region of the HSV-1 UL15 gene consists of a first exon and a second exon separated by a single intron. UL15 mRNA is generated by splicing reactions that delete the central intron (9, 10, 11). At the non-permissive temperature (NPT), the UL15 temperature-sensitive (ts) mutant was able to replicate the viral genome but was unable to package the viral genome into the capsid (12). Furthermore, concatemeric viral genomes accumulated in the UL15 ts mutant-infected cells at the NPT (13). Roizman et al. reported that in HSV-1 (F)-infected cells, two types of UL15 proteins [with molecular weights (Mw) of 35 kDa and 75 kDa] shared the C-terminus of the second exon of UL15 (13). The 75 kDa UL15 protein was required for viral genome cleavage and packaging, and HSV-1 mutants with a genetic disruption of UL15 failed to form the mature C-capsids and instead formed the immature B-capsids (14). Since a 35 kDa protein is detected in HSV-1 mutant-infected cells with stop codons inserted into the first exon of UL15, this indicated that the 35 kDa protein is the translation product of the second exon of UL15 (15). The translation product from the second exon of UL15 was referred to as UL15.5. UL15.5 was not required for HSV-1 infectious virus production, but the function of UL15.5 is still unknown (16). Baines et al. also reported that full-length UL15 and the two low Mw forms of UL15 (3 kDa-and 4 kDa-reduced forms) were detected in purified B-capsids (17, 18). Taken together, these findings indicated that UL15 plays a role in both viral genome cleavage and packaging into the capsid. Moreover, the UL15 gene expresses a variety of proteins with different Mws.

The KSHV ORF29 gene encodes a 687 amino acid (aa) protein. This gene contains one intron between the first and second exons (19, 20). Le Grice et al. found that the ORF29 C-terminal domain has DNA sequence-independent nuclease activity (21). We demonstrated that ORF29 interacts with ORF7 but not with ORF67.5 (6). Moreover, the interaction between ORF7 and ORF67.5 is enhanced by ORF29, and the interaction between ORF29 and ORF7 is enhanced by ORF67.5 (8). Glaunsinger et al. generated an ORF29.stop virus in which Leu338 and Gln339 of KSHV ORF29 are changed to stop codons. The production of infectious virus generated from the ORF29.stop mutant was impaired, and expression of the L gene (K8.1) was decreased at the mRNA and protein levels. In addition, viral genome replication was also reduced in ORF29.stop KSHV (22). Although several functions of ORF29 were reported, the contribution of ORF29 to KSHV terminase function remains unresolved.

In this paper, we have attempted to characterize ORF29 by generating KSHV lacking full-length ORF29 (ORF29-deficient KSHV) and its revertant. ORF29-deficient KSHV was impaired in virus production and infectious virion production, but interestingly, neither K8.1 expression nor viral genome replication was impaired. ORF29-deficient KSHV failed to form mature capsids and instead formed an immature capsid, the soccer ball-like capsid. It also lost the TR cleavage ability. In addition, we detected a novel KSHV protein (designated ORF29.5), which was expressed in the lytic phase. Furthermore, we found that ORF29 preferentially interacted with itself rather than with ORF7.

## Results

### Construction of an ORF29-deficient KSHV bacterial artificial chromosome (BAC)

To analyze the function of KSHV ORF29, we constructed a full-length ORF29-deficient KSHV BAC clone (ΔORF29-BAC16) and its reverse-mutated KSHV BAC clone (Revertant-BAC16). We generated ΔORF29-BAC16 by introducing a frameshift via deleting a C-G bp located 1 bp downstream of the ORF29 start codon (ATG) (Fig. 1A). Although ORF34 and ORF34.1 are present near the ORF29 start codon in the KSHV genome, the mutation introduction site in ΔORF29-BAC16 does not overlap with the ORF34 or ORF34.1 coding region. The generated ΔORF29-BAC16 clone contains nonsense mutations in the ORF29 region. Thus, the ΔORF29-BAC16 clone has DNA sequences encoding 25 aa unrelated to ORF29 from the start codon to 75 bp within ORF29 followed by a stop codon at 76-78 bp. Furthermore, Revertant-BAC16 was generated by reinsertion of the missing 1 bp (i.e., C-G bp) into ΔORF29-BAC16 (Fig. 1A). The nucleotide sequences around the mutagenesis sites of ΔORF29-BAC16 and Revertant-BAC16 were confirmed by Sanger sequencing (Fig. 1A). Each BAC clone was digested with the restriction enzyme EcoRI, and the insertion and removal of the kanamycin resistance gene during mutagenesis was confirmed by changes in signal patterns obtained by agarose gel electrophoresis (Fig. 1B). Wild-type (WT)-BAC16, ΔORF29-BAC16, and Revertant-BAC16 were transfected into iSLK cells and selected with hygromycin B to establish BAC16 stably harboring cells, which were defined as WT-iSLK cells, ΔORF29-iSLK cells, and Revertant-iSLK cells, respectively. When iSLK cells harboring BAC16 (i.e., KSHV latently-infected cells) are treated with doxycycline (Dox) and sodium butyrate (SB), the lytic infection is effectively induced via Dox-elicited RTA expression (23). Next, we attempted to confirm the loss of ORF29 expression in ΔORF29-iSLK cells and the return of ORF29 expression in Revertant-iSLK cells by Western blotting (WB) using an antibody (Ab) specific for ORF29. A rabbit anti-ORF29 polyclonal antibody (pAb) was raised using the 22 to 41 aa region (GERWELSAPTFTRHCPKTAR) within ORF29 as a peptide antigen (Fig. 1C). Each cell line was treated (or untreated) with Dox and SB for 72 h to induce lytic infection. We attempted to detect endogenous ORF29 by WB but were unable to detect ORF29 due to numerous non-specific signals (data not shown). Therefore, the ORF29 protein was immunoprecipitated with anti-ORF29 pAb and the precipitates were subjected to WB. The calculated Mw of endogenous ORF29 is 76.5 kDa. When lytic infection was induced, ORF29 expression was detected in WT-iSLK and Revertant-iSLK cells, but not in ΔORF29-iSLK cells (Fig. 1D). In all cell lines, ORF29 was not detected in the non-lytic state (Fig. 1D).

**FIG 1.**
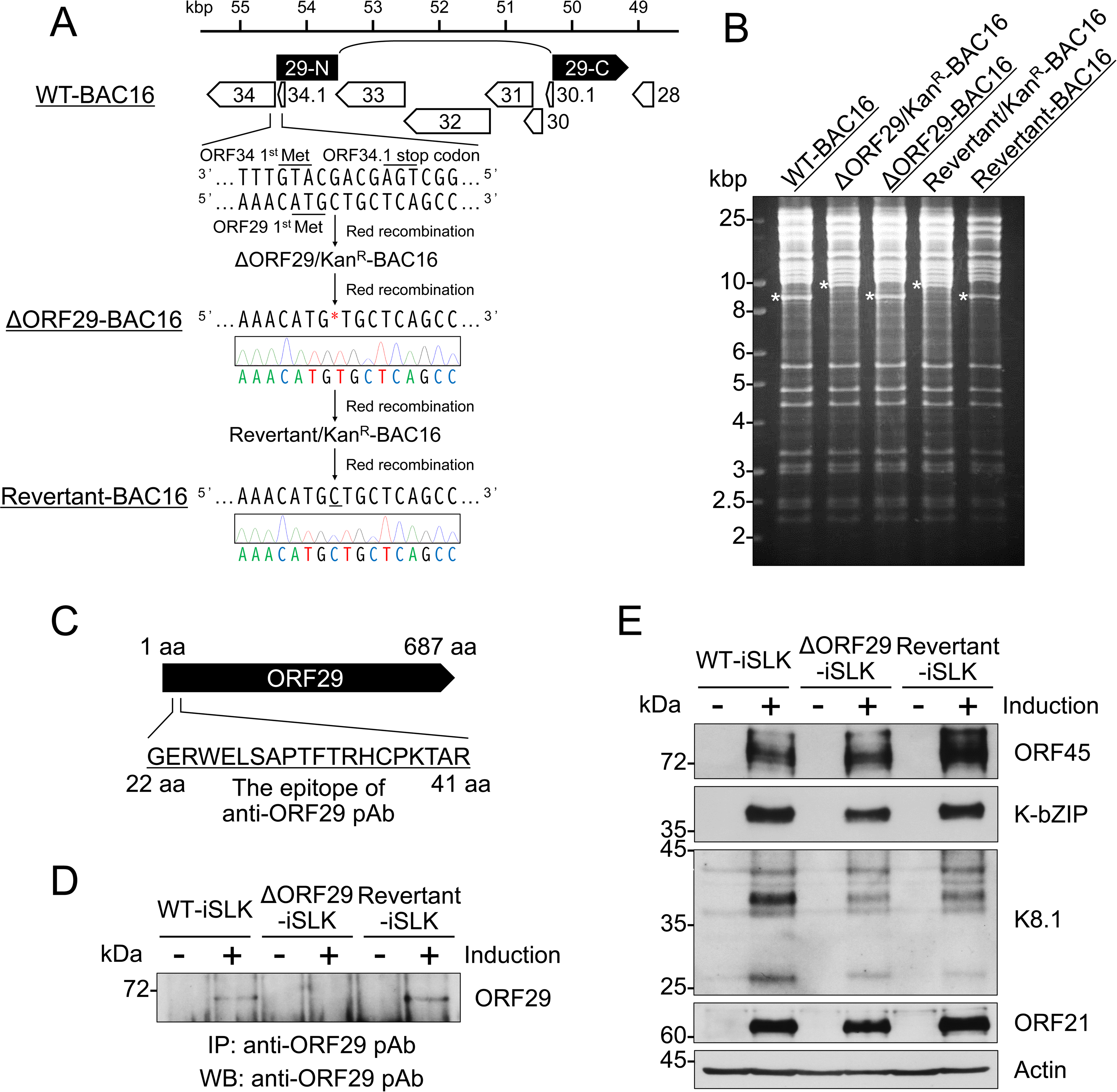
Construction of ΔORF29-BAC16 and Revertant-BAC16. (A) Conceptual diagram showing the location of KSHV ORF29 (nucleotides 49179 to 50321 and 53572 to 54492; accession number: GQ994935) and the process for generating each mutant KSHV-BAC. The nucleotide sequence adjacent to each mutagenesis site was determined by Sanger sequencing. (B) Each BAC DNA was digested with EcoRI, and the DNA segments were separated by agarose gel electrophoresis to confirm the insertion and removal of the kanamycin resistance cassette. The asterisks indicate the insertion or deletion of the kanamycin resistance cassette in each BAC clone. (C) The antigenic peptide sequence for preparation of the rabbit anti-ORF29 pAb; aa, amino acids. (D) Validation of endogenous ORF29 protein expression in lytic-induced (+) or uninduced (−) WT-iSLK cells, ΔORF29-iSLK cells, and Revertant-iSLK cells. WT-BAC16, ΔORF29-BAC16, and Revertant-BAC16 were transfected into iSLK cells, and stably BAC16-harboring iSLK cells were established (defined as WT-iSLK cells, ΔORF29-iSLK cells, and Revertant-iSLK cells, respectively). Each cell line was treated (or untreated) with Dox and SB for 72 h to induce lytic replication. Next, cell lysates were subjected to IP with anti-ORF29 pAb-binding Dynabeads conjugated to Protein G. The immunoprecipitates were subjected to WB with anti-ORF29 pAb. (E) Protein expression of lytic genes in lytic-induced (+) or uninduced (−) WT-iSLK cells, ΔORF29-iSLK cells, and Revertant-iSLK cells. Each cell line was treated (or untreated) with Dox and SB for 72 h and subjected to WB with the respective antibodies. Actin was used as a loading control.

In addition to ORF29, the expression of other lytic proteins in each iSLK cell line were examined. The expression of the IE protein ORF45, the DE protein K-bZIP, the L protein K8.1, and ORF21 were detected in WT-iSLK cells, ΔORF29-iSLK cells, and Revertant-iSLK cells under the lytic phase (Fig. 1E). These results indicated that a deficiency in either the ORF29 protein or the ORF29 gene in lytic-induced KSHV harboring cells does not remarkably affect the expression of ORF45, K-bZIP, K8.1, and ORF21 proteins.

### Viral genome replication and virion production are impaired in ORF29-deficient KSHV

To assess the contribution of ORF29 to virus production, we evaluated the transcription of lytic genes, the amount of intracellular viral genome replication, and virus production in lytic-induced ΔORF29-iSLK cells. WT-iSLK cells, ΔORF29-iSLK cells, and Revertant-iSLK cells were treated with Dox and SB for 72 h, and the cells and culture supernatants were collected. The mRNA expression levels of the IE gene, ORF16, the DE gene, ORF46 + ORF47, and the L gene, K8.1, were measured by reverse transcription-quantitative PCR (RT-qPCR) using total RNA from harvested cells. There were no remarkable differences in the expression levels of the tested lytic genes in each cell line (Fig. 2A). Next, the intracellular viral genome copy number from harvested cells was quantified by qPCR. The amount of intracellular viral genome was comparable in each cell line (Fig. 2B). Next, to evaluate the extracellular encapsidated viral genome, copy number, DNase-treated culture supernatants were quantified by qPCR. We found that the encapsidated viral copy number derived from the ΔORF29-iSLK cell line supernatant was significantly decreased compared to that of WT-iSLK and Revertant-iSLK cell lines (Fig. 2C). Next, infectious virus production from ΔORF29-iSLK cells was examined by a supernatant transfer assay. The culture supernatants from each lytic-induced iSLK cell line were co-cultured with fresh HEK293T cells, and the GFP positivity of the HEK293T cells was measured by flow cytometry. Since BAC16 contains the GFP gene, the infectivity of BAC16-derived KSHV can be assessed by measuring GFP-derived fluorescence (24). The amount of infectious virus produced by ΔORF29-iSLK cells was markedly reduced compared with the amount of infectious virus produced by WT-iSLK and Revertant-iSLK cells (Fig. 2D). These results indicated that ORF29 is not essential for KSHV lytic gene expression or viral genome replication, but is important for the production of infectious virions.

**FIG 2.**
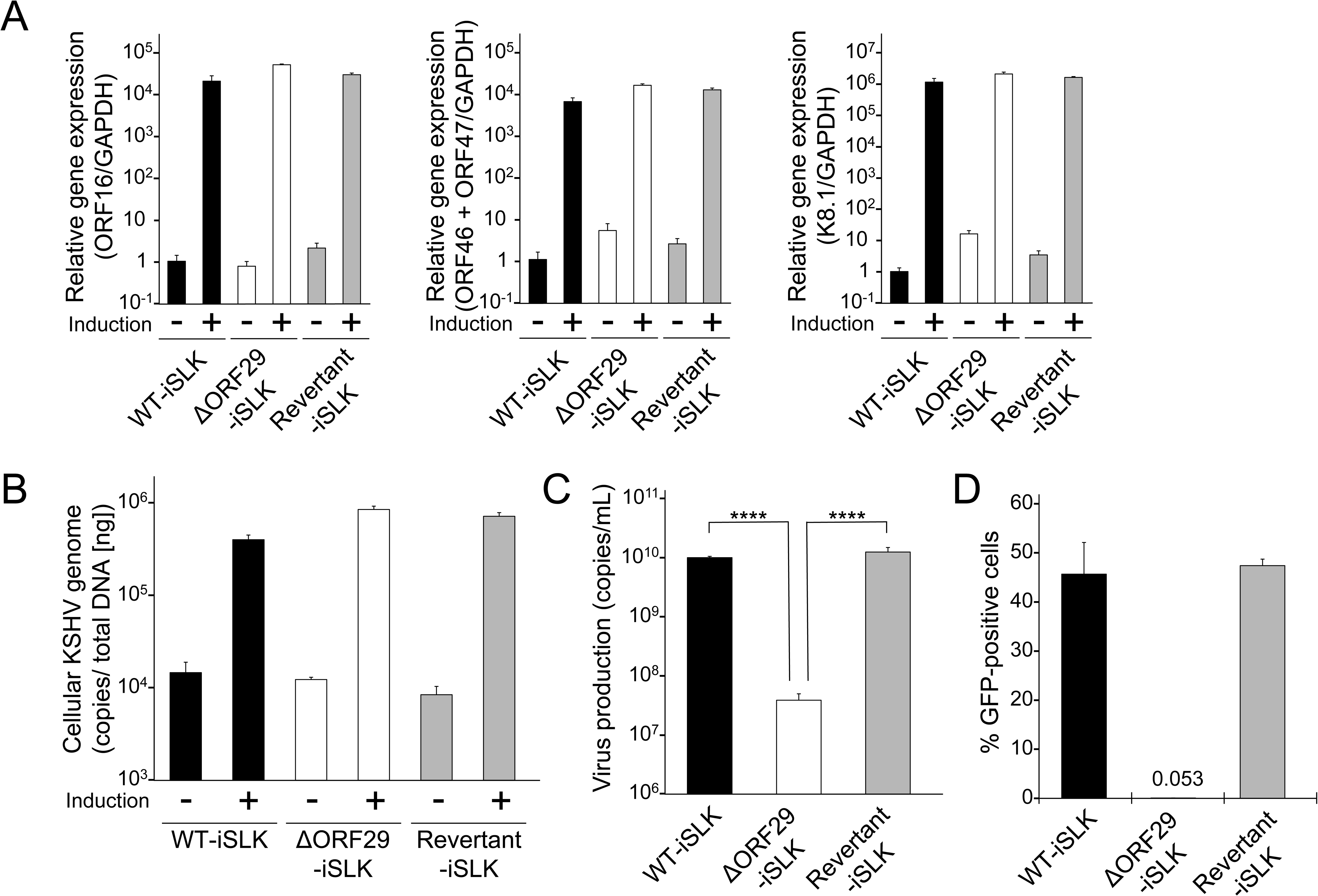
Characterization of ΔORF29 KSHV. (A) Transcription of KSHV lytic genes in lytic-induced (+) or uninduced (−) WT-iSLK cells, ΔORF29-iSLK cells, and Revertant-iSLK cells. Each cell line was treated (or untreated) with Dox and SB for 72 h to induce lytic replication. Total RNA was extracted from the cells and subjected to RT-qPCR to evaluate mRNA expression of the following lytic genes: IE gene (ORF16), DE gene (ORF46 and ORF47), and L gene (K8.1). The viral gene mRNA levels were normalized to GAPDH mRNA levels. The values obtained from uninduced WT-iSLK cells were defined as 1.0. (B) Intracellular KSHV genome copy number. Each cell line was treated (or untreated) with Dox and SB, and the intracellular DNA was extracted. The KSHV genome copy number was quantified by qPCR. The genome copy number was normalized to the total DNA content. (C) Extracellular encapsidated KSHV genome copy number. The cell lines were treated with Dox and SB for 72 h, and the culture supernatant was collected. The encapsidated viral genome was purified from the DNase-treated supernatant, and the viral genome copy number was quantified by qPCR. ****, P < 0.001. (D) Production of infectious virions. Each cell line was treated with Dox and SB for 72 h, and the culture supernatant was analyzed with a supernatant transfer assay. The culture supernatants were used for infection of fresh HEK293T cells. At 24 h post-infection, the percentage of GFP-positive HEK293T cells was measured by flow cytometry to evaluate infectious virus production.

To further examine the contribution of ORF29 to virion production, we performed a complementation assay for virion production in ΔORF29-iSLK cells. The extracellular encapsidated viral genome copy number was reduced in ΔORF29-iSLK cells compared to WT-iSLK cells. However, this reduction was significantly restored by transient transfection of a C-terminal FLAG-tagged ORF29 (ORF29-FLAG) expression plasmid (Fig. 3A). The expression of exogenous ORF29-FLAG was confirmed by WB (Fig. 3B). Furthermore, the ability of ΔORF29-iSLK cells to produce infectious virus, which was inhibited due to the loss of ORF29, was rescued by transfection of the ORF29-FLAG expression plasmid (Fig. 3C). These results confirmed that ORF29 is required for virion production.

**FIG 3.**
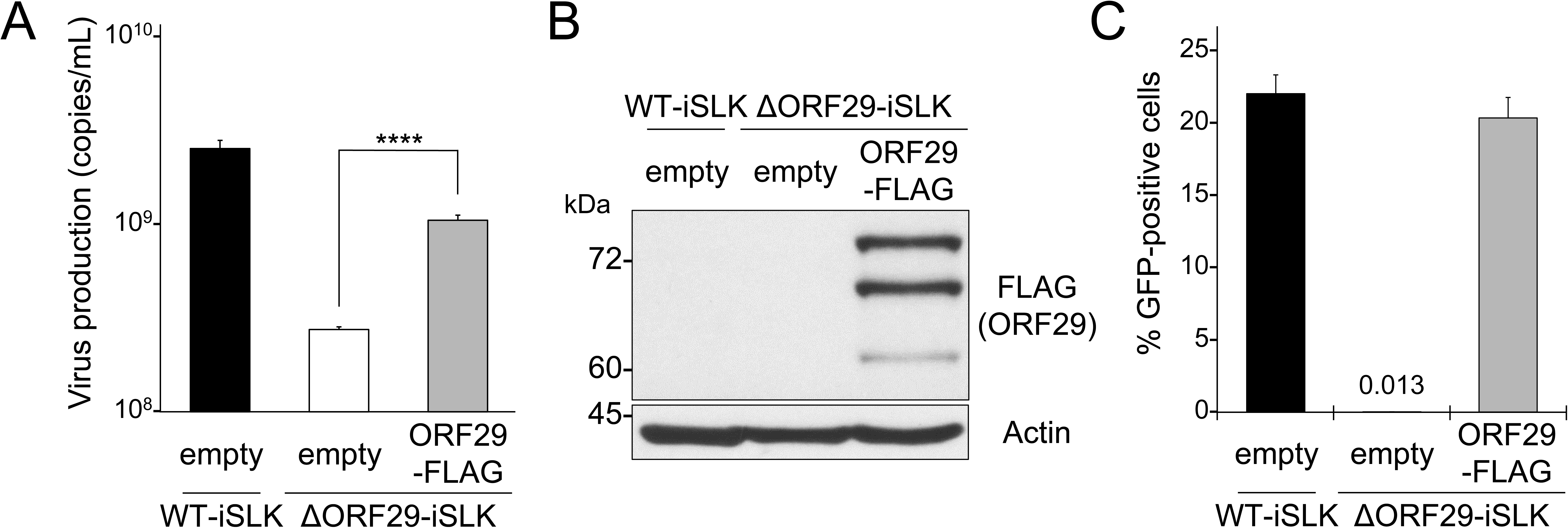
Complementation of reduced virion production in ORF29-deficient KSHV by exogenous ORF29 expression. (A) Transient expression of exogenous ORF29 complemented the reduction in extracellular encapsidated viral genomes detected in ΔORF29-iSLK cells. ΔORF29-iSLK cells were transiently transfected with the ORF29-FLAG plasmid or with the control plasmid lacking the ORF29 gene (empty). Next, the cells were treated with Dox and SB for 72 h to induce lytic replication. Encapsidated viral genomes in the culture supernatant was quantified by qPCR. ****, P < 0.001. (B) Exogenous ORF29 expression in the samples described in Fig. 3A was confirmed by WB using anti-FLAG Ab. (C) Transient expression of exogenous ORF29 rescued the decrease in infectious virion production detected in ΔORF29-iSLK cells. ΔORF29-iSLK cells were transiently transfected with the ORF29-FLAG plasmid or with the control plasmid lacking the ORF29 gene (empty). Next, the cells were treated with Dox and SB for 72 h. The harvested culture supernatants were analyzed with a supernatant transfer assay. HEK293T cells were infected with produced virions in the culture supernatant. At 24 h post-infection, the percentage of GFP-positive HEK293T cells was measured by flow cytometry.

### ORF29-deficient KSHV capsid maturation is arrested at the immature soccer ball-like capsid stage

The following three types of KSHV capsid structures have been defined from electron microscopic images: A-capsid, which is an empty capsid; B-capsid, which contains a globular scaffold protein but lacks the viral genome; and C-capsid, which is a mature capsid containing the viral genome but lacks the scaffold protein (25). We have previously reported that KSHV deficient in ORF7 or ORF67.5, which are components of the KSHV terminase complex, are unable to form mature capsids and form immature soccer ball-like capsids (7, 8). To investigate the contribution of ORF29 to capsid maturation, the morphology of capsids formed in WT-iSLK and ΔORF29-iSLK cell lines were observed by transmission electron microscopy (TEM). Cells were treated with Dox and SB for 48 h, and the nuclear capsids were photographed. In WT-iSLK cells, A-capsids, B-capsids, C-capsids, and soccer ball-like capsids were detected (Fig. 4A). In contrast, C-capsids were not observed in ΔORF29-iSLK cells, and most of the capsids formed were soccer ball-like capsids (Fig. 4B). Fig. 4C shows the number of capsids of each type. The data indicated that the KSHV capsid formation process was arrested at the immature soccer ball-like capsid stage in lytic-induced ΔORF29-iSLK cells. Thus, similar to ORF7 and ORF67.5, ORF29 was also important for KSHV capsid maturation.

**FIG 4.**
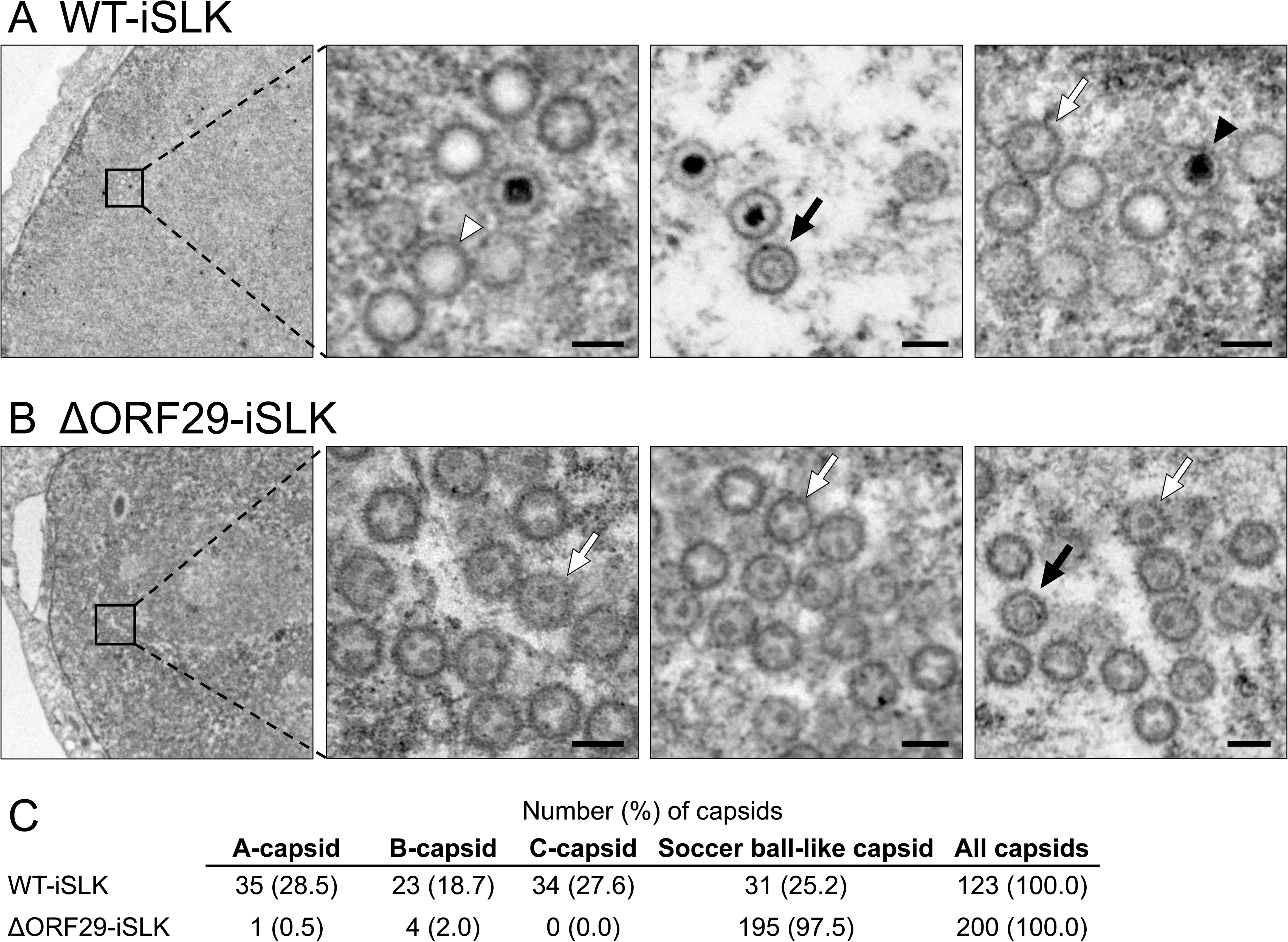
The soccer ball-like capsids were mainly produced in lytic-induced ΔORF29-iSLK cells. TEM images showed the morphology of capsids formed in the nuclei of (A) WT-iSLK cells and (B) ΔORF29-iSLK cells during the lytic phase. The cells were treated with Dox and SB for 48 h to induce the lytic phase, and the nuclear capsids were observed by TEM. The black arrowheads, white arrowheads, black arrows, and white arrows indicate C-capsids, A-capsids, B-capsids, and soccer ball-like capsids, respectively. Scale bars, 100 nm. (C) Quantification of each type of capsid observed in lytic-induced WT-iSLK and ΔORF29-iSLK cells.

### ORF29 is essential for cleavage of the TRs in the KSHV genome

The terminase complex encapsidates the single unit-length viral genome into the capsid and cleaves the TR of the viral genome precursor, resulting in the formation of a mature capsid. Proper cleavage at the TR site within the viral genome precursor by the terminase complex can be assessed by Southern blotting using a probe directed against 1xTR (7, 8, 26). To analyze the contribution of ORF29 to the terminase complex genome cleavage activity, TR cleavage within the viral genome precursor was compared in ΔORF29-iSLK and WT-iSLK cells. Total DNA prepared from lytic-induced iSLK cells was digested with EcoRI and SalI restriction enzymes to release the TRs, and the TRs were detected by Southern blotting. Cleaved TRs were not detected in WT-iSLK cells, ΔORF29-iSLK cells, and Revertant-iSLK cells under the non-lytic state. When lytic infection was induced, the cleaved TRs were detected in WT-iSLK and Revertant-iSLK cells, but not in ΔORF29-iSLK cells (Fig. 5). These results indicated that ORF29 is required for the TR cleavage activity of the KSHV terminase complex. In addition, the amount of uncleaved TRs was increased by lytic induction in all cell lines (Fig. 5). This result is consistent with the results of Fig. 2B showing that ΔORF29-iSLK cells undergo viral genome replication.

**FIG 5.**
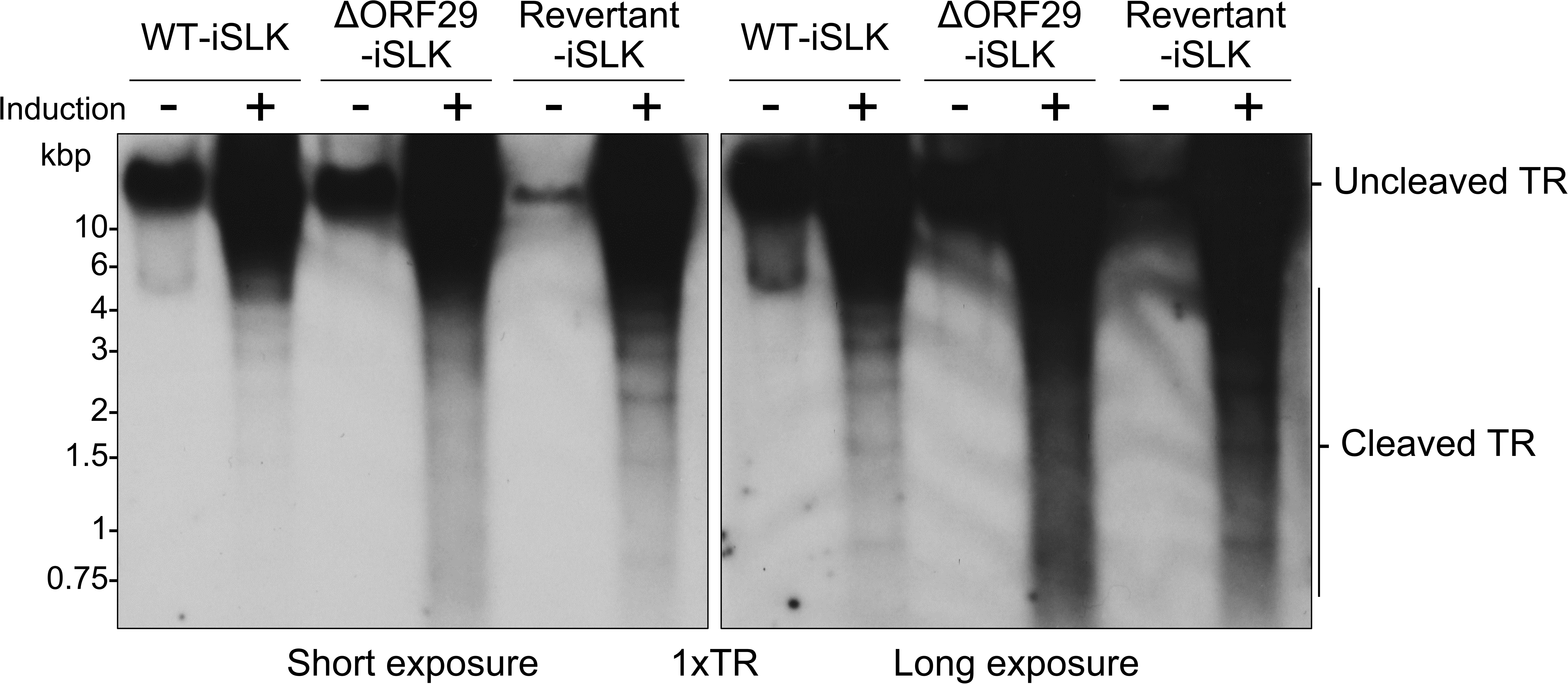
ORF29-deficient KSHV failed to cleave the TRs in the viral genome precursor. WT-iSLK cells, ΔORF29-iSLK cells, and Revertant-iSLK cells were treated (or untreated) with Dox and SB for 72 h to induce lytic replication. The intracellular genomic DNA was purified from lytic-induced (+) or uninduced (−) cells. The genomic DNA was digested with EcoRI and SalI, and the digested DNA was subjected to Southern blotting. Uncleaved and cleaved TRs from the KSHV genome were detected using the digoxigenin (DIG)-labeled 1xTR probe; left, short exposure; right, long exposure.

### The N-terminal region of ORF29 is required for its interaction with ORF7, and full-length ORF29 is required for enhancing the formation of the terminase complex

The KSHV terminase complex is thought to be comprised of ORF7, ORF29, and ORF67.5. ORF7, ORF29, and ORF67.5 form a tripartite complex, but ORF29 and ORF67.5 do not interact directly (6). ORF7 interacts with both ORF29 and ORF67.5 and serves as the hub molecule of the tripartite complex (6). To elucidate the responsible region in ORF29 which is required for its interaction with ORF7, plasmids encoding ORF29 partial deletions (Δ1: Δ2-100 aa, Δ2: Δ101-200 aa, Δ3: Δ201-300 aa, Δ4: Δ301-400 aa, Δ5: Δ301-400 aa, Δ6: Δ501-600 aa, Δ7: Δ601-687 aa) were generated (Fig. 6A). A pulldown assay was performed to analyze whether each of the ORF29 deletion mutants possessed the ability to interact with ORF7. The C-terminal S-tagged ORF7 (ORF7-S) plasmid and each C-terminal FLAG-tagged ORF29 mutant (ORF29-FLAG) plasmid were co-transfected into HEK293T cells. Next, the ORF7-S protein was pulled down (Pd) from the cell lysates using agarose beads immobilized to the S-protein which binds to the S-tag. ORF29 WT and Δ2-7 interacted with ORF7, however, the interaction between ORF29 Δ1 and ORF7 was not detected (Fig. 6B). These results suggested that the N-terminal region of ORF29 (specifically aa 2-100) is important for the interaction between ORF29 and ORF7.

**FIG 6.**
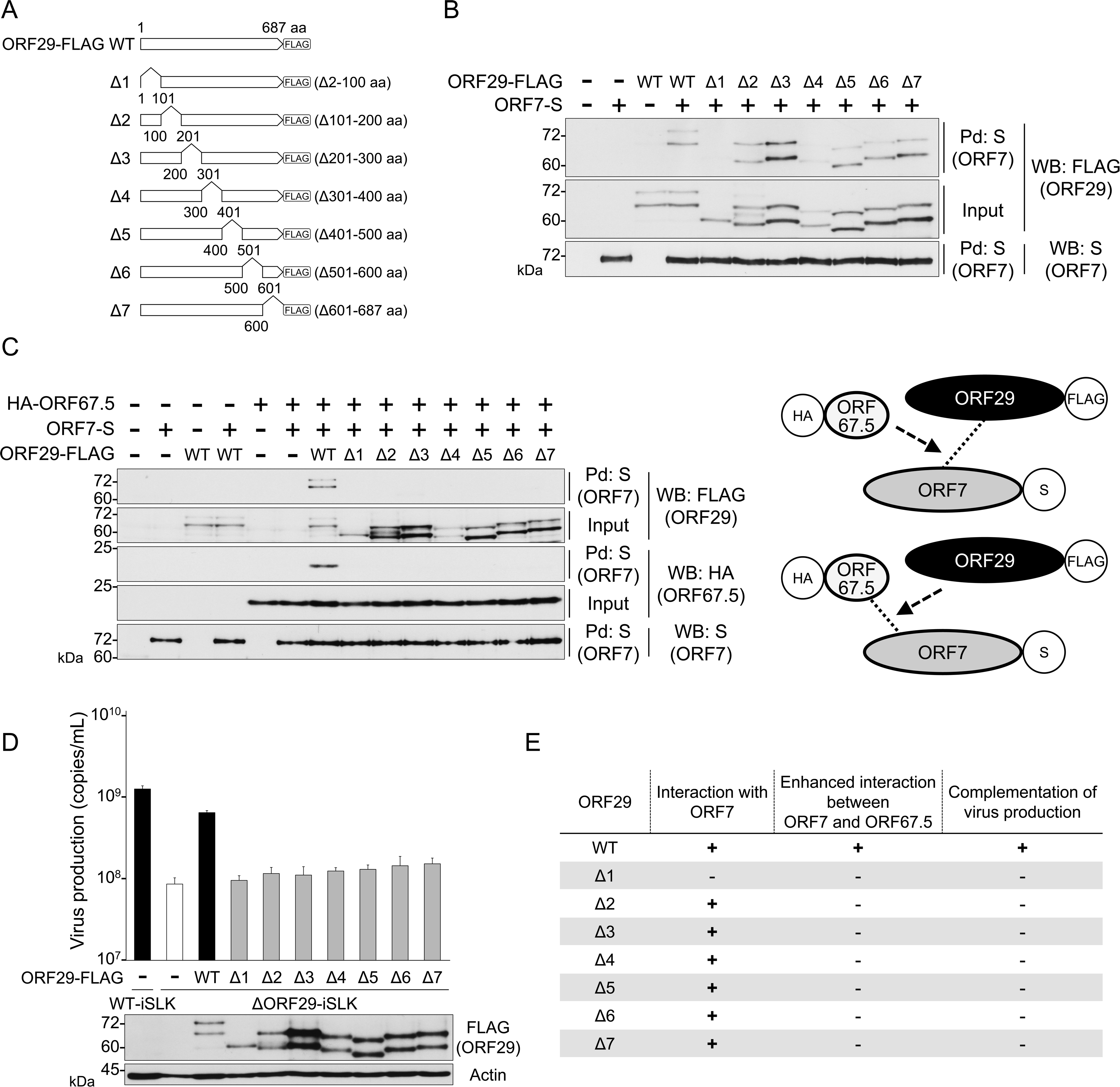
The N-terminal region (aa 2-100) of ORF29 is required for its interaction with ORF7. (A) Schematic representation of the C-terminal FLAG-tagged ORF29 deletion mutants. The deleted aa are shown to the right of the corresponding mutant. (B) The N-terminal aa 2-100 of ORF29 was essential for its interaction with ORF7. The interaction of ORF29 deletion mutants (Δ1-Δ7) with WT ORF7 was analyzed by pulldown assays. WT ORF7-S plasmid (ORF7-S) and ORF29-FLAG deletion mutant plasmids were co-transfected into HEK293T cells. The cells were lysed in lysis buffer, and ORF7-S protein was Pd using S-protein agarose. The precipitates were subjected to WB. (C) All ORF29 deletion mutants lost the ability to enhance the interaction between ORF7 and ORF67.5. Plasmids encoding HA-ORF67.5, ORF7-S, and ORF29-FLAG deletion mutants were co-transfected into HEK293T cells, and the cells were lysed in lysis buffer. The ORF7-S protein was Pd by S-protein agarose, and the interaction partner of the precipitated ORF7-S protein was detected by WB. The right side shows a schematic representation of the tested interaction models. (D) Complementation of the reduction in encapsidated viral genome production in ORF29-deficient KSHV cells by ORF29 deletion mutants. ΔORF29-iSLK cells were transfected with the respective ORF29 mutant plasmids and were cultured in medium with Dox and SB for 72 h to induce the lytic phase. The culture supernatants were collected, and the encapsidated viral genome copy number in the culture supernatant was quantified by qPCR. The bottom panel shows a WB using an anti-FLAG Ab to confirm the expression of exogenous ORF29 mutant proteins. (E) A summary of the results obtained in Figs. 6B, C, and D.

We reported that the interaction between ORF7 and ORF67.5 is enhanced by ORF29, and the interaction between ORF7 and ORF29 is enhanced by ORF67.5 (8). Therefore, we investigated the ability of the ORF29 deletion mutants to enhance the interaction between ORF7 and ORF67.5. The ORF7-S plasmid, the N-terminal HA-tagged ORF67.5 (HA-ORF67.5) plasmid and each mutated ORF29-FLAG plasmid were co-transfected into HEK293T cells. Next, the ORF7-S protein was Pd with S-protein agarose. As expected, the interaction between ORF7 and ORF67.5 was enhanced by ORF29 WT, and the interaction between ORF7 and ORF29 WT was also enhanced by ORF67.5 (Fig. 6C). In contrast to WT ORF29, all ORF29 deletion mutants (i.e., ORF29 Δ1-7) failed to enhance the interaction between ORF7 and ORF67.5. Moreover, ORF67.5 did not enhance the interaction between ORF7 and all ORF29 Δ1-7 mutants (Fig. 6C). The results showed that only full-length ORF29 retained the ability to enhance the interaction between the components of the terminase complex. The right panel of Fig. 6C shows a schematic of the interaction of each ORF evaluated in this experiment.

Finally, the rescue of virus production by the ORF29 deletion mutants was evaluated with a complementation assay. ΔORF29-iSLK cells were transfected with each ORF29 mutant plasmid, and the DNase-resistant viral genome copies in the culture supernatant were quantified. Virus production in ΔORF29-iSLK cells was reduced compared to WT-iSLK cells, however, this reduction was rescued by overexpression of ORF29 WT in ΔORF29-iSLK cells (black bars in Fig. 6D). In contrast, overexpression of all ORF29 deletion mutants (ORF29 Δ1-7) failed to restore virus production (gray bars in Fig. 6D). These results indicated that full-length ORF29 is required for virus production. The characteristics of ORF29 WT and Δ1-7 are summarized in Fig. 6E. The results indicated that aa 2-100 of ORF29 are required for ORF29 to interact with ORF7. Moreover, full-length ORF29 is important for the enhanced interaction between the KSHV terminase complex components and for the virus-producing function of ORF29. However, we should consider the possibility that these ORF29 mutants do not retain the native conformation of WT ORF29.

### Exogenous expression of the ORF29 plasmid revealed that translation is initiated not only from the 1^st^ AUG, but also from the 2^nd^, 3^rd^, and 4^th^ AUGs in the ORF29 mRNA

The transient transfection of the C-terminal tagged ORF29 plasmid into iSLK cells expressed multiple forms of ORF29 with different Mws (Fig. 3B). To analyze this phenomenon, we examined the expression patterns of ORF29 produced from the N-terminal S-tagged ORF29 (S-ORF29) and the C-terminal S-tagged ORF29 (ORF29-S) plasmids (Fig. S1A). Each plasmid was transfected into HEK293T cells, and S-tagged ORF29 proteins were Pd with S-protein agarose and were subjected to WB with anti-S Ab. Based on the estimated Mw, one signal derived from the full-length ORF29 was detected for S-ORF29. In contrast, multiple signals derived from low Mw ORF29 were detected for ORF29-S in addition to the signal derived from full-length ORF29 (Fig. S1A). From these results, we hypothesized that ORF29 undergoes proteolytic cleavage near its N-terminal region. To test this hypothesis, ORF29-S proteins of lower Mw than full-length ORF29-S were subjected to N-terminal aa sequence analysis by Edman degradation, but the analysis was not possible (data not shown). This suggested that the low Mw proteins are not protease cleavage products and instead are acetylated at the first Met. Another possibility is that these short forms are the translation products initiated at the 1^st^, 2^nd^, 3^rd^, and 4^th^ AUG of the ORF29 mRNA transcribed from the transfected ORF29-S plasmid. Therefore, we constructed C-terminal FLAG-tagged point mutant ORF29 plasmids (ORF29-FLAG [M42V, M105V, M173V]) in which the 2^nd^, 3^rd^, and 4^th^ Met codons were replaced with Val codons (Fig. S1B). Lysates from HEK293T cells transfected with each plasmid were subjected to immunoprecipitation (IP) with anti-FLAG Ab. The estimated Mws of the full-length ORF29 and the translation products from the point mutant ORF29 plasmids (including the FLAG tag portion) are as follows: full-length, 79.7 kDa; M42, 75.0 kDa; M105, 68.0 kDa; and M173, 60.2 kDa. Full-length ORF29 was detected in cells transfected with ORF29 WT, M42V, M105V, or M173V plasmids (Fig. S1B). Meanwhile, in ORF29 M42V, M105V, and M173V plasmid-transfected cells, we observed the disappearance of signals whose Mw corresponded to the low Mw ORF29 detected in the ORF29-FLAG WT-expressing cells (Fig. S1B). In addition, we constructed C-terminal FLAG-tagged deletion mutant ORF29 plasmids (ORF29-FLAG [M42, M105, M173]) whose translation is initiated at the 2^nd^, 3^rd^, and 4^th^ Met codons in ORF29 (Fig. S1C). The Mw of each signal appearing in ORF29 M42, M105, and M173 plasmid-expressing cells was completely consistent with the low Mw signals of ORF29 appearing in ORF29 WT plasmid-expressing cells (Fig. S1C). These data indicated that exogenous expression of ORF29 by transfection with our plasmid expresses not only the full-length ORF29 translation product, but also translation products from the 2^nd^, 3^rd^, and 4^th^ Met codons in the ORF29 mRNA.

### The ORF29 short forms exogenously expressed by transfection with the ORF29 plasmid do not contribute to ORF29 function in terminase complex formation and in virus production

The results of Fig. S1 raised the possibility that the ORF29 short forms translated from the 2^nd^, 3^rd^, and 4^th^ AUGs of the ORF29 mRNA affected the results of the plasmid-based assay. We tested whether ORF29 short forms expressed by plasmid transfection could play a role in the properties and functions of ORF29, i.e., interaction with ORF7, enhancement of the interaction of ORF7 with ORF67.5, and recovery of the decrease in virus production in ΔORF29-iSLK cells. The interaction of ORF7 with the ORF29 deletion mutants produced by the ORF29 M42, M105, or M173 plasmid used in Fig. S1C was confirmed by pulldown assays. The interaction of ORF29 WT with ORF7 was detected, and ORF29 M42 also interacted with ORF7. However, neither ORF29 M105 nor M173 interacted with ORF7 (Fig. S2A). Regarding the ability to enhance the interaction between ORF7 and ORF67.5, ORF29 WT and M42, but not ORF29 M105 and M173, had this ability (Fig. S2B). Finally, the contribution of ORF29 short forms to virus production was examined. The reduction in virus production in ΔORF29-iSLK cells was restored by transfection of the ORF29-WT plasmid, but not by transfection of the ORF29 M42, M105, or M173 plasmids (Fig. S2C). These data indicated that ORF29 M42 retains the ability to interact with ORF7 and enhances the interaction between ORF7 and ORF67.5. However, full-length ORF29 translated from the 1^st^ AUG is required for ORF29 function in virus production.

### The KSHV genome encodes a novel lytic gene, ORF29.5

Since exogenous expression by transfection with the ORF29 plasmid expressed multiple forms of ORF29 translated from the 2^nd^, 3^rd^, and 4^th^ Met codons (Figs. 3B, S1, and S2), we analyzed whether these phenomena also occurred with endogenous ORF29 expression. Since the anti-ORF29 pAb recognizes the N-terminal region of ORF29, this pAb cannot detect the short forms of ORF29 lacking the N-terminal side (Figs. 1C and D). Therefore, we attempted to establish a rabbit pAb using a peptide antigen consisting of the 646-662 aa region of ORF29 (RDGGQSYSAKQKHMSDD), which comprises the C-terminal region within the 687 aa full-length ORF29. The generated Ab failed to detect specific signals of ORF29 (data not shown). Next, we generated a modified KSHV-BAC with a FLAG-tag DNA sequence inserted at the 3’ end of the ORF29 coding region to detect the endogenous expression of FLAG-conjugated ORF29 protein by WB. We then generated the new modified WT-BAC16 and ΔORF29-BAC16 clones in which the FLAG-tag-coding DNA sequence was inserted just before the stop codon of the ORF29 gene. These modified BAC clones were named WT-29CtermF-BAC16 and ΔORF29-29CtermF-BAC16 (Figs. 7A and B). We expected that the WT-29CtermF-BAC16 clone will produce the C-terminal FLAG-tagged ORF29 (ORF29-FLAG) protein, whereas the ΔORF29-29CtermF-BAC16 clone will not. Next, WT-29CtermF-BAC16 or ΔORF29-29CtermF-BAC16 was transfected into iSLK cells and selected with hygromycin B to establish BAC16 stably harboring cells, which were defined as WT-29CtermF-iSLK orΔORF29-29CtermF-iSLK, respectively. WT-iSLK cells, ΔORF29-iSLK cells, WT-29CtermF-iSLK cells, and ΔORF29-29CtermF-iSLK cells were treated with Dox and SB for 72 h. Newly synthesized ORF29-FLAG proteins were IPd with FLAG-tag Ab-immobilized beads, and the precipitated ORF29 was detected by WB using FLAG-tag Ab. Surprisingly, in addition to the full-length ORF29-FLAG protein (72 kDa), a novel 30 kDa translated product with a FLAG-tag was detected in the lytic-induced WT-29CtermF-iSLK cells. Meanwhile, only a 30 kDa translated product with a FLAG-tag was detected in the lytic-induced ΔORF29-29CtermF-iSLK cells (Fig. 7C). Thus, the mRNA encoding a 30 kDa protein detected in both WT-29CtermF-iSLK cells and ΔORF29-29CtermF-iSLK cells possess the same stop codon and FLAG-tag DNA sequence as the mRNA encoding the full-length ORF29 protein. Here, this new translation product with a Mw of 30 kDa was referred to as ORF29.5 (Fig. 7C). We did not detect the short forms of ORF29 (75, 68, and 60.2 kDa) translated from the 2^nd^, 3^rd^, and 4^th^ Met codons of the ORF29 mRNA observed in Fig. S1 (Fig. 7C). No FLAG-positive signals were detected in all samples under the non-lytic state (Fig. 7C). In summary, KSHV expresses two translation products during lytic infection: the 72 kDa full-length ORF29 and the 30 kDa ORF29.5.

**FIG 7.**
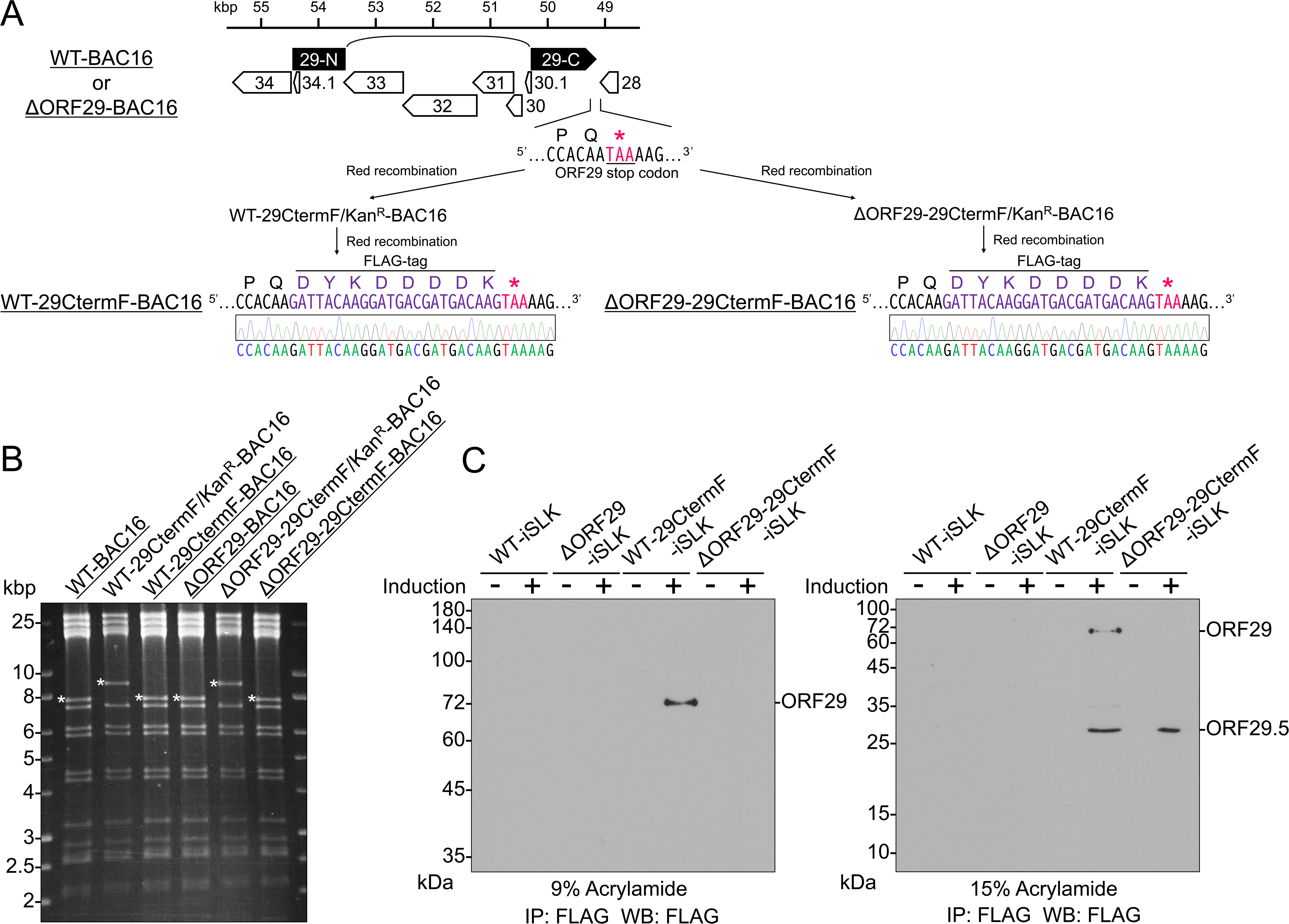
KSHV expresses a novel viral protein, ORF29.5, in the lytic phase. (A) Diagram showing the modified KSHV-BACs with a FLAG-tag DNA sequence inserted at the 3’ end of the ORF29 coding region. To detect the endogenous expression of ORF29 protein by WB using anti-FLAG Ab, a FLAG-tag DNA sequence was inserted before the stop codon of the ORF29 gene in WT-BAC16 and ΔORF29-BAC16 clones. These modified clones were defined as WT-29CtermF-BAC16 and ΔORF29-29CtermF-BAC16. (B) Each BAC clone was digested with SalI and agarose gel electrophoresis was used to confirm the insertion and removal of the kanamycin resistance cassette. The asterisks indicate the insertion or deletion of the kanamycin resistance cassette in each BAC clone. (C) Expression of a novel viral protein, ORF29.5, in lytic-induced WT-29CtermF-iSLK and ΔORF29-29CtermF-iSLK cells, which have a FLAG-tag DNA sequence inserted at the 3’ end of the ORF29 coding region. WT-29CtermF-BAC16 or ΔORF29-29CtermF-BAC16 was transfected into iSLK cells, and stably BAC16-harboring iSLK cells were established (defined as WT-29CtermF-iSLK or ΔORF29-29CtermF-iSLK, respectively). Each cell line was treated (induction +) or untreated (induction -) with Dox and SB for 72 h to induce the lytic phase. The cells were then lysed in RIPA buffer and endogenously expressed FLAG fusion proteins were precipitated with anti-FLAG Ab-immobilized beads. The precipitates were subjected to WB from 9% (left) or 15% (right) polyacrylamide gels.

### ORF29 is a self-interacting protein

The KSHV terminase complex is comprised of ORF7, ORF29, and ORF67.5. The heteromeric protein-protein interactions of the terminase complex components have been elucidated. Specifically, it has been shown that ORF7 interacts with both ORF29 and ORF67.5, but ORF29 and ORF67.5 do not interact with each other (6). Previously, it was unknown whether the terminase complex proteins self-interact. Therefore, we examined whether ORF7 is a self-interacting protein. The ORF7-FLAG plasmid and either the ORF7-S or ORF29-S plasmid were co-transfected into HEK293T cells, and the ORF7-S protein or ORF29-S protein was Pd with S-protein agarose. The precipitated molecule (i.e., the interacting partner of ORF7-S or ORF29-S) was probed by WB. As expected, the interaction between ORF29-S and ORF7-FLAG was detected. However, we did not detect a self-interaction of ORF7-S-FLAG (Fig. 8A). Next, the self-interaction of ORF29 was examined. The ORF29-FLAG plasmid and either the ORF29-S or ORF7-S plasmid were co-transfected into HEK293T cells, and ORF29-S or ORF7-S protein was Pd to detect its binding partner. As expected, the interaction between ORF7-S and ORF29-FLAG was detected. Interestingly, ORF29-FLAG interacted more strongly with ORF29-S than with ORF7-S (Fig. 8B). Finally, the self-interaction of ORF67.5 was evaluated. The HA-ORF67.5 plasmid and either the S-ORF67.5 or ORF7-S plasmid were co-transfected into HEK293T cells, and the S-ORF67.5 or ORF7-S protein was Pd with S-protein agarose. A weaker self-interaction was detected for ORF67.5 compared to the interaction with ORF7 (Fig. 8C). These results demonstrated that ORF29 interacted more strongly with itself than with ORF7, i.e., ORF29 was found to be a self-interacting protein.

**FIG 8.**
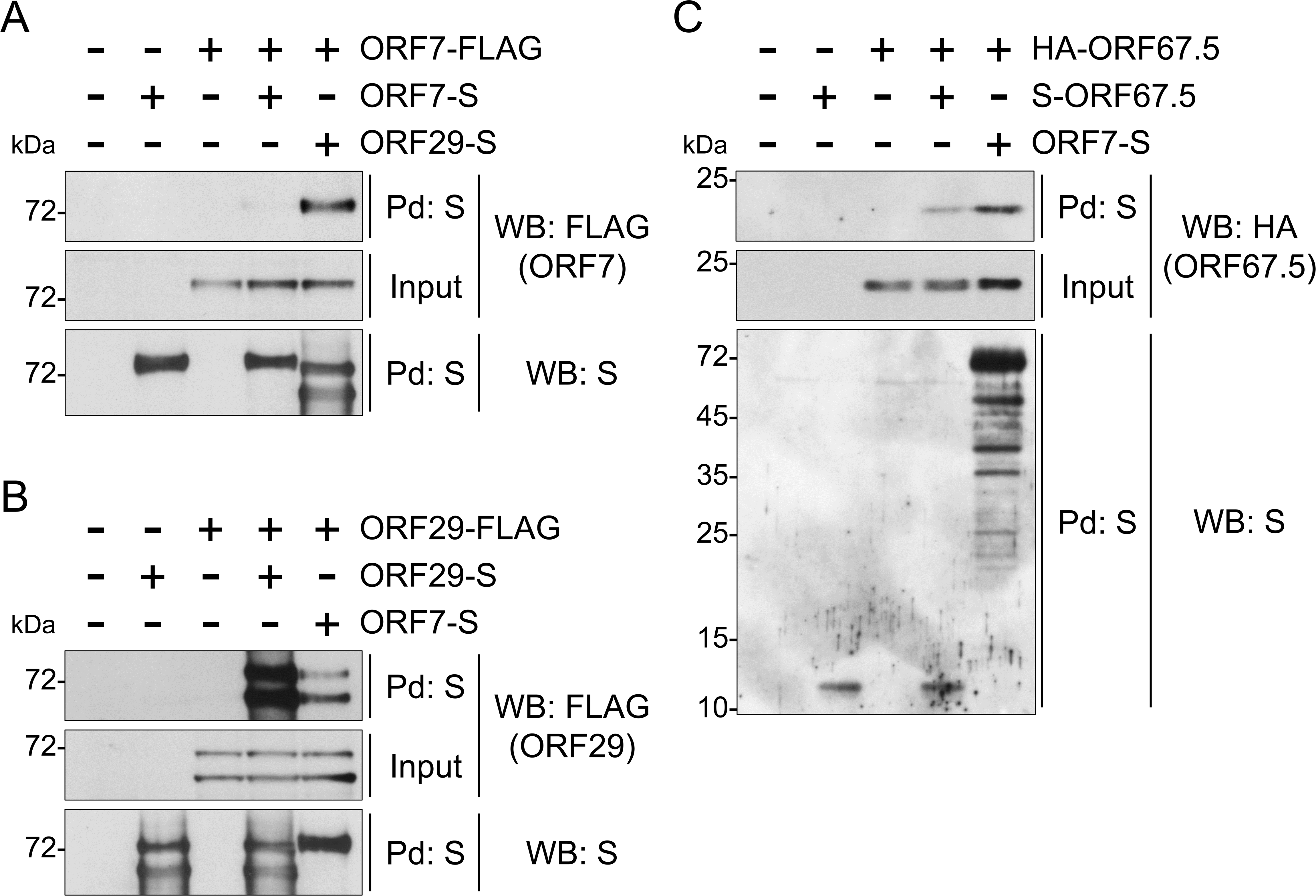
ORF29 preferentially interacted with itself rather than with ORF7. The self-interaction activities of each component of the KSHV terminase complex were evaluated by pulldown assays. Plasmids were co-transfected into HEK293T cells, and the cells were lysed in lysis buffer. (A, B) ORF29-S, (A, B, C) ORF7-S or (C) S-ORF67.5 protein was Pd with S-protein agarose, and the precipitated proteins were analyzed by WB using an anti-FLAG-tag Ab for detection of (A, B) ORF7 and ORF29, or an anti-HA-tag Ab for detection of (C) ORF67.5.

## Discussion

In this study, we showed that KSHV ORF29 is a component of the terminase complex and plays essential roles in both TR cleavage of viral genome precursors and capsid maturation. In addition, full-length ORF29 was required for both enhancement of terminase complex formation and virus production. Moreover, we showed that in the lytic phase, KSHV expresses a 30 kDa ORF29.5 that shares the same stop codon of ORF29. We also found that ORF29 is a self-interacting protein. To our knowledge, this is the first report demonstrating that ORF29 contributes to terminase function as a component of the KSHV terminase complex. This finding was established by characterization of a fully ORF29-deficient KSHV. In addition, there have been no reports regarding the novel KSHV lytic gene product that we refer to as ORF29.5.

ORF29-deficient KSHV failed to process the viral genome precursor and predominantly formed the soccer ball-like capsids in the nuclei of infected cells (Figs. 4B, 4C, and 5). Similarly, ORF7-or ORF67.5-deficient KSHVs fail to process viral genome precursors and predominantly form soccer ball-like capsids (7, 8). Our previous studies and the results of this paper indicated that when KSHV terminase function is impaired, capsid maturation is arrested at an immature capsid stage. The soccer ball-like capsids produced by ORF7-, ORF29-, and ORF67.5-deficient KSHVs contain a common internal structure resembling the telstar pattern or coin dot of a soccer ball (7, 8). It has been hypothesized that when the KSHV genome is encapsidated into the capsid, the decayed scaffold proteins are extruded and eliminated from the capsid (25). We speculate that the soccer ball-like capsids are immature capsids in which decayed scaffold proteins are not shed from the capsid but remain within the capsid (7). As our data show, the loss of KSHV terminase function (i.e., failure to process the viral genome precursor) resulted in the formation of soccer ball-like capsids. This fact supports the hypothesis presented in the literature (25) that decayed scaffold proteins are ejected from the capsid by genome packaging.

The N-terminal region of ORF29 was shown to be important for its interaction with ORF7 (Fig. 6B). An ORF29 deletion mutant lacking the aa 2-100 region showed no interaction with ORF7 (Fig. 6B). In contrast, the ORF29 deletion mutant lacking aa 1-41 interacted with ORF7 (Fig. S2A). Based on these results, the 42-100 aa region of ORF29 is important for its interaction with ORF7. In addition to the interaction with ORF7, the 42-100 aa region of ORF29 may be important for the formation of the proper conformation of ORF29. However, each deletion mutant of ORF29 that was found to interact with ORF7 was also unable to enhance the interaction between ORF7 and ORF67.5 (Figs. 6B and C). Based on these results, we hypothesized that each deletion mutant of ORF29 may not be able to form a terminase complex. Taken together, these results suggested that the entire ORF29 protein is required for the formation of the proper conformation of the terminase complex.

We detected a novel KSHV lytic-expressed 30 kDa protein referred to as ORF29.5. During the lytic state, ORF29 expression was not detected in ΔORF29-29CtermF-iSLK cells, but ORF29.5 was expressed in ΔORF29-29CtermF-iSLK and WT-29CtermF-iSLK cells (Fig. 7C). These results indicated that ORF29.5 mRNA uses the same stop codon of ORF29 mRNA. Based on these data, we hypothesize that at least part of the coding region in the ORF29.5 mRNA overlaps with the 3’-terminal coding region of the ORF29 mRNA. We also speculate that ORF29.5 is expressed in the lytic-induced ΔORF29-iSLK cells used in this study. Since ORF29-deficient KSHV failed to cleave the TR in the KSHV genome (Fig. 5), the deficiency of ORF29 resulted in a loss of terminase function even in the presence of ORF29.5 expression. This indicated that ORF29.5 cannot complement the function of ORF29 as a component of the KSHV terminase complex. The virological function and the mechanism of ORF29.5 expression are currently unknown. In addition, during the lytic phase, ORF29-deficient KSHV (ΔORF29-29CtermF-BAC16) expressed ORF29.5 despite not expressing ORF29 (Fig. 7C), which indicated that ORF29 expression is not required for ORF29.5 expression. Furthermore, our data indicated that the ORF29.5 protein is not derived from ORF29 but is its own translation product. ORF29-deficient KSHV (ΔORF29-29CtermF-BAC16) lacks 1 bp immediately after the ORF29 start codon, suggesting that ORF29.5 may begin translation at the AUG codon downstream of the ORF29 start codon. However, the translation start site of ORF29.5 is currently unknown.

Based on previous studies, we discuss the ORF29.5 expression mechanism. The first exon of the mRNA precursor encoding ORF29 is annotated as ORF29a, and the second exon as ORF29b. The second exon contains the stop codon for ORF29 (3). The 3’ end of ORF29a contains a splice donor site and the 5’ end of ORF29b contains a splice acceptor site. The splicing reaction links the 3’ end of ORF29a to the 5’ end of ORF29b, resulting in the production of mature ORF29 mRNA (19, 20). We propose the following three possible mechanisms for ORF29.5 expression. First of all, translation is initiated from a novel translation start site located between the ORF29 1^st^ Met codon and the stop codon in ORF29 mRNA, resulting in the expression of ORF29.5, a shorter form than ORF29. Second, ORF29.5 is translated from the ORF29b mRNA, which is fused to mRNAs encoding other ORFs. It was reported that ORF48-ORF29b chimeric mRNA, in which an ORF48 mRNA and an ORF29b mRNA are linked, was generated by splicing reactions and was expressed as an IE transcript (27). The ORF48 protein is a negative regulator of the cGAS/STING pathway (28). In addition to ORF48-ORF29b, ORF40-29b chimeric mRNA, in which ORF40 mRNA and ORF29b mRNA are linked, was also reported to be generated by splicing reactions (29). The ORF40 gene encodes the N-terminal aa region of the primase-associated factor involved in viral genome replication (30, 31). There is a putative intron of 19,656 nucleotides between ORF29b and ORF48 and a putative intron of 10,195 nucleotides between ORF29b and ORF40 (29). It is not known what proteins are translated from ORF48-ORF29b mRNA and ORF40-ORF29b mRNA, but it is possible that ORF29.5 is expressed from these mRNAs. The final possible mechanism of ORF29.5 expression is that the ORF29.5 protein may be translated from the mRNA transcribed from the transcription start site within the ORF29b encoding DNA. This is due to the presence of a putative TATA-like promoter (TTATTAAAAA) within the DNA sequence of ORF29b (32). As for the end of translation, the ORF29.5 mRNA and ORF29 mRNA are thought to share a common stop codon located in the ORF29 coding region (Fig. 7C). The virological and physiological characterization of ORF29.5, including its mechanism of expression, needs to be elucidated by further studies.

In a previous study of ORF29, Glaunsinger et al. generated the ORF29.stop virus expressing the N-terminal region of ORF29. Their results showed that ORF29 is important for the L gene expression and KSHV genome replication (22). However, characterization of our ΔORF29-BAC16 clone generated in this paper did not reveal any impairment of K8.1 (L gene) expression nor KSHV genome replication due to ORF29 deletion (Figs. 1E, 2A, 2B, and 5). One possible explanation for the difference between their results and our results may be the difference of the introduced mutation site in the ORF29 gene. Our ΔORF29-BAC16 has a frameshift mutation induced by deleting a C-G bp located 1 bp downstream of the ORF29 start codon. This mutation generates a nonsense ORF29 mRNA, which has a stop codon 76-78 bp downstream of the ORF29 start codon and is destined for degradation. In fact, ΔORF29-BAC16 did not express ORF29 protein (Figs. 1A and D). On the other hand, the ORF29.stop virus used in the previous study was generated by changing the 338^th^ and 339^th^ codons of ORF29 to stop codons (22). Therefore, it is assumed that the ORF29.stop virus expresses the N-terminal region (aa 1-337) of the full-length ORF29 protein (687 aa). This C-terminally truncated ORF29 protein may inhibit the L gene expression and KSHV genome replication.

The conformation of the KSHV terminase complex has not yet been determined, but the conformation of the HSV-1 terminase complex has been determined by cryo-electron microscopy (33). The components of the HSV-1 terminase complex are UL15, UL28, and UL33, which are homologs of KSHV ORF29, ORF7, and ORF67.5, respectively. UL28 interacts with UL15 and UL33 (34, 35). One molecule each of UL15, UL28, and UL33 forms a tripartite complex, and six of these tripartite complexes assemble into a ring to form a hexameric ring (33). The HSV-1 genome passes within this terminase ring, and genome packaging into capsids occurs (33). In this study, KSHV ORF29 was found to be a robustly self-interacting protein (Fig. 8B), while KSHV ORF67.5 exhibited a weaker self-interaction. The self-interaction of ORF29 (also ORF67.5) may contribute to the multimerization of a terminase complex or the formation of the terminase ring via self-assembly of multiple ORF29 molecules. The significance of the self-interacting feature of ORF29 is unknown and needs to be clarified in the future.

We have focused on the KSHV terminase complex and have characterized its components. Currently, ORF7, ORF29, and ORF67.5 are considered as components of the KSHV terminase complex. Our previous studies have shown that ORF7 and ORF67.5 are important for KSHV terminase function (6, 7, 8). This study showed that ORF29 is also important for KSHV terminase function. Thus, we have made progress in the functional characterization of the KSHV terminase complex. The herpesvirus terminase machinery is essential for viral replication, but it is not present in humans. Therefore, the KSHV terminase complex is one of the most promising targets for anti-KSHV drug development. In addition, elucidation of the currently unresolved conformation of the KSHV terminase complex will promote both further understanding of this complex and the development of anti-KSHV drugs.

## materials and methods

### Cell culture and reagents

HEK293T cells were cultured in Dulbecco’s modified Eagle’s medium (DMEM) (Nacalai Tesque Inc., Kyoto, Japan) supplemented with 10% fetal bovine serum (FBS). iSLK cells (23) were cultured in DMEM supplemented with 10% FBS, 1 μg/mL of puromycin (InvivoGen, CA, USA), and 0.25 mg/mL of G418 (Nacalai Tesque, Inc.). iSLK cells harboring KSHV-BAC were cultured in DMEM supplemented with 10% FBS, 1 μg/mL of puromycin (InvivoGen), 0.25 mg/mL of G418 (Nacalai Tesque, Inc.), and 1 mg/mL of hygromycin B (Wako, Osaka, Japan).

### Plasmids

The N-terminal 2xS-tagged ORF29 expression plasmid (YI-31), the C-terminal 3xFLAG-tagged ORF29 expression plasmid (RF-009), the C-terminal 2xS-tagged ORF7 expression plasmid (YI-02), the C-terminal 3xFLAG-tagged ORF7 expression plasmid (YI-04), the N-terminal 2xS-tagged ORF67.5 expression plasmid (YI-16), and the N-terminal 5xHA-tagged ORF67.5 expression plasmid (YI-17) have been described previously (6, 8). The C-terminal 2xS-tagged ORF29 expression plasmid (RF-010) was constructed by PCR using the previously constructed N-terminal 3xFLAG-tagged ORF29 expression plasmid (YI-52) as a template and digesting the obtained insert with EcoRI (Takara Bio, Shiga, Japan) and SalI (TOYOBO, Osaka, Japan) (6). The DNA ligation kit Mighty Mix (Takara Bio) was used for ligation. Each C-terminal 3xFLAG-tagged ORF29 mutant expression plasmid (Δ1 [YI-94], Δ2 [YI-95], Δ3 [YI-96], Δ4 [YI-97], Δ5 [YI-98], Δ6 [YI-99], Δ7 [YI-100], M42V [YI-101], M105V [YI-102], M173V [YI-103], M42 [YI-105], M105 [YI-106], and M173 [YI-107]) was constructed using the In-Fusion HD Cloning kit (Takara Bio). The inserts were obtained by PCR using the C-terminal 3xFLAG-tagged ORF29 expression plasmid (RF-009) as a template. The KOD-Plus-Neo (TOYOBO) was used for PCR, and the pCI-neo mammalian expression vector (Promega, WI, USA) was used as the backbone vector. The primers used for plasmid construction are listed in Table 1. The sequences of the inserts were confirmed by Sanger sequencing.

**TABLE 1.**
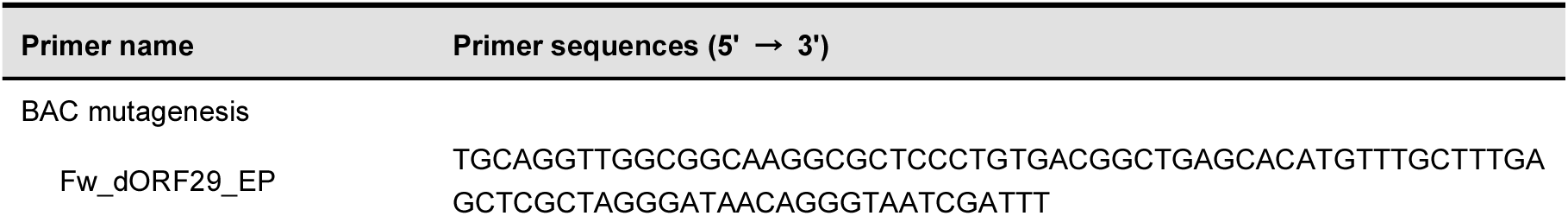
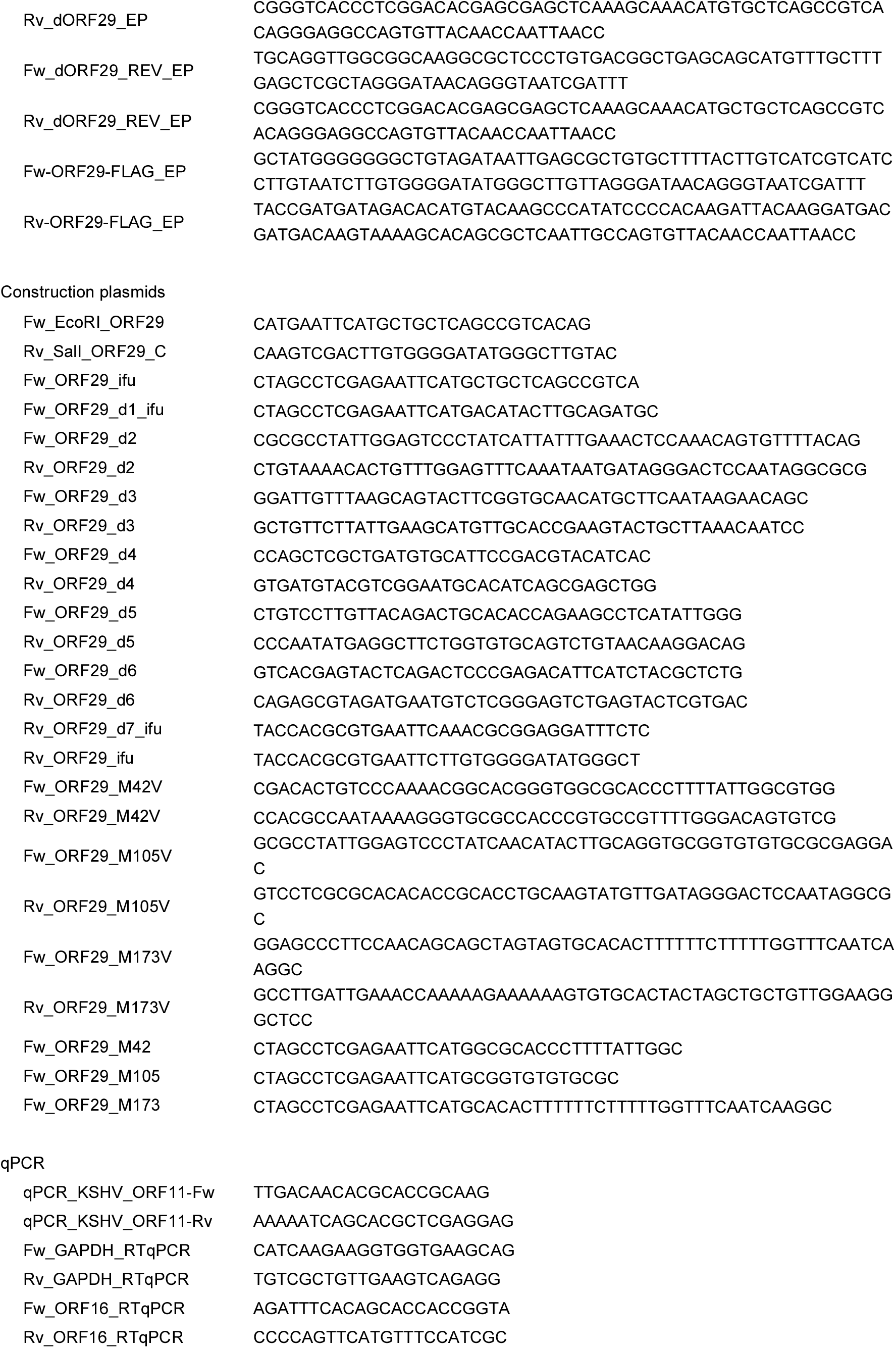
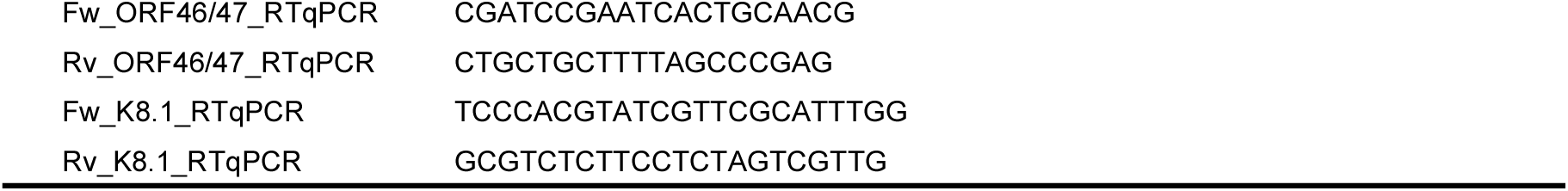
Primers for BAC mutagenesis, construction of plasmids, and qPCR.

### Mutagenesis of BAC16 clones

Mutagenesis of BAC16 clones were performed as described in previous publications (24, 36). ΔORF29-BAC16 was constructed by deleting a single G at position 54,489 in WT-BAC16 (accession number: GQ994935). Revertant-BAC16 was generated by reinsertion of the G missing in ΔORF29-BAC16. WT-29CtermF-BAC16 and ΔORF29-29CtermF-BAC16 were constructed by inserting a FLAG-tag sequence between the Ala at position 49,181 and the Tyr at position 49,182 in WT-BAC16 and ΔORF29-BAC16 (accession number: GQ994935). The primers used for this mutagenesis are shown in Table 1. The insertion and deletion of kanamycin resistance cassettes in each BAC16 clone were analyzed by EcoRI or SalI digestion and agarose gel electrophoresis. The mutated sites of each BAC16 clone were verified by Sanger sequencing.

### Generation of iSLK cells stably harboring individual BAC16 clones

Each BAC16 clone was purified from the Escherichia coli strain GS1783 by NucleoBond Xtra BAC (Takara Bio). WT-BAC16 or its mutants (ΔORF29-BAC16, Revertant-BAC16, WT-29CtermF-BAC16, and ΔORF29-29CtermF-BAC16) were transfected into iSLK cells by the calcium phosphate method. The transfected iSLK cells were selected under 1 mg/mL of hygromycin B (Wako) to establish Dox-inducible recombinant KSHV stably harboring cell lines (WT-iSLK, ΔORF29-iSLK, Revertant-iSLK, WT-29CtermF-iSLK, and ΔORF29-29CtermF-iSLK).

### Measurement of viral gene expression, intracellular viral genome replication, and virus production

Each measurement was performed according to previously described methods with slight modifications (8). Briefly, iSLK cells harboring WT or each BAC16 mutant were treated with 8 μg/mL of Dox and 1.5 mM of SB for 72 h to induce lytic replication.

To measure viral gene expression, lytic-induced or uninduced cells (3.5 × 10^5^ cells in a 6-well plate) were harvested with 500 μL of RNAiso Plus (Takara Bio). The extracted total RNA was treated with DNase I (New England Biolabs, MA, USA) and resuspended in 300 uL of RNAiso Plus (Takara Bio). The cDNA was synthesized from 160 ng of DNase-treated total RNA using ReverTra Ace qPCR RT Master Mix (TOYOBO). qPCR was performed using the synthesized cDNA as a template with the THUNDERBIRD Next SYBR qPCR mix (TOYOBO). Table 1 lists the primers used to measure viral gene expression. The relative mRNA expression levels were determined by the delta-delta threshold cycle (ΔΔCT) method and were normalized to GAPDH mRNA levels.

To quantify intracellular viral genome replication, iSLK cells (3.5 × 10^4^ cells in a 48-well plate) were induced or uninduced and harvested. Viral genome DNA and cellular genomic DNA were purified from the harvested cells using a QIAamp DNA Blood mini kit (Qiagen, CA, USA). The viral genome copy number was quantified by qPCR and normalized to the amount of total DNA. qPCR assays were performed using the THUNDERBIRD Next SYBR qPCR mix (TOYOBO) and the KSHV ORF11-specific primers, which are listed in Table 1.

To quantify extracellular encapsidated viral DNA, iSLK cells (1.5 × 10^5^ cells in a 12-well plate) were induced, and culture supernatants were harvested and centrifuged to remove debris. The supernatants were treated with DNase I (New England Biolabs), and encapsidated viral DNA was extracted from the supernatants using a QIAamp DNA Blood mini kit (Qiagen). Purified viral DNA copy numbers were quantified by qPCR using the THUNDERBIRD Next SYBR qPCR mix (TOYOBO) and the KSHV ORF11-specific primers.

Infectious virus production was quantitated using a supernatant transfer assay. iSLK cells (2 × 10^6^ cells on a 10 cm dish) were induced, and the culture supernatants and cells were collected. The supernatants and cells were centrifuged, and the supernatants were mixed with fresh HEK293T cells (7.5 × 10^5^ cells) and polybrene (8 μg/mL; Sigma-Aldrich, MO, USA). The mixtures were added to 12-well plates. The plates were centrifuged at 1,200 *g* for 2 h and incubated for 24 h. GFP-positive cells were detected with a flow cytometer (FACSCalibur, Beckton Dickinson, CA, USA) using CellQuest Pro software (Beckton Dickinson).

### Complementation assay

iSLK cells (1.5 × 10^5^ cells in a 12-well plate for quantification of extracellular encapsidated viral DNA, or 2 × 10^6^ cells on a 10 cm dish for measurement of infectious virus production) were induced with 8 μg/mL of Dox and 1.5 mM of SB and concurrently transfected with each plasmid. After 72 h, the respective evaluations were conducted according to the methods described in the section entitled Measurement of viral gene expression, intracellular viral genome replication, and virus production.

### Western blotting

Cells were washed with phosphate-buffered saline (PBS) and lysed in SDS sample buffer [50 mM Tris-HCl (pH 6.8), 5% (wt/vol) SDS, 50% (vol/vol) glycerol, 0.002% (wt/vol) bromophenol blue, and 2% (vol/vol) 2-mercaptoethanol]. Next, the samples were sonicated for 10 s, reduced at 60°C for 20 min, and subjected to SDS-PAGE. The ExcelBand All Blue Regular Range Protein Marker (PM1500) (SMOBIO, Hsinchu County, Taiwan) was used as a Mw marker. The proteins were transferred to a ClearTrans nitrocellulose membrane 0.2 μm (Wako), and the membrane was incubated for 30 min at room temperature in 5% (wt/vol) nonfat dry milk in PBS with 0.1% (vol/vol) Tween-20 (PBS-T). The membrane was then incubated with a primary Ab followed by incubation with a secondary Ab in Can Get Signal Immunoreaction Enhancer Solution (TOYOBO). Immunodetection was achieved with the ECL Western Blotting Detection Reagents (Cytiva, Tokyo, Japan). The blot was then exposed to X-ray film (Fuji film, Tokyo, Japan).

### Antibodies

Anti-KSHV ORF29 rabbit pAb was generated by GL Biochem, Shanghai, China, using the synthetic peptide GERWELSAPTFTRHCPKTAR (ORF29: aa 22 to 41) as the antigen. Anti-KSHV ORF29 rabbit pAb was purified from the immunized rabbit serum using antigen peptide affinity chromatography. The following primary antibodies were used: anti-ORF45 mouse monoclonal Ab (mAb; 2D4A5; Santa Cruz Biotechnology, TX, USA), anti-K-bZIP mouse mAb (F33P1; Santa Cruz Biotechnology), anti-K8.1 A/B mouse mAb (4A4; Santa Cruz Biotechnology), anti-ORF21 rabbit pAb (previously produced in our laboratory) (37), anti-beta actin mouse mAb (AC-15; Santa Cruz Biotechnology), anti-FLAG-tag mouse mAb (FLA-1; MBL, Nagoya, Japan), anti-S-tag rabbit pAb (sc-802; Santa Cruz Biotechnology), and anti-HA-tag mouse mAb (TANA2; MBL). Anti-mouse IgG-horseradish peroxidase-conjugated (HRP; NA931; Cytiva) and anti-rabbit IgG-HRP (7074; Cell Signaling Technology, MA, USA) were used as the secondary antibodies. HRP conjugated anti-FLAG-tag mouse mAb (M2; Sigma-Aldrich, MO, USA) was used in Figs. S1B, S1C, and 7C.

### Immunoprecipitation and pulldown assay

Ten 10 cm dishes of each iSLK cell line harboring BAC (2 × 10^6^ cells/dish) were induced or uninduced for 72 h. HEK293T cells (2 × 10^6^ cells on a 10 cm dish) were transfected with 12 μg of plasmid DNA and 36 μg of polyethylenimine hydrochloride (PEI) MAX (Polysciences, Inc., PA, USA) for 20 h. The cells were lysed in lysis buffer [50 mM Tris-HCl (pH 8.0), 120 mM NaCl, 1% (vol/vol) glycerol, 0.2% (vol/vol) Nonidet P-40 substitute, and 1 mM dithiothreitol] or RIPA buffer [50 mM Tris-HCl (pH 8.0), 150 mM NaCl, 1% (vol/vol) Nonidet P-40 substitute, 0.5% (wt/vol) sodium deoxycholate, and 0.1% (wt/vol) SDS]. The buffers used for lysis are described in each figure legend. The cell extracts were incubated with appropriate beads for 1 h, and the beads were washed three times. S-protein agarose (Merck KGaA, Darmstadt, Germany), Dynabeads Protein G (Thermo Fisher Scientific, MA, USA), and Anti DYKDDDDK tag Antibody Magnetic Beads (1E6; Wako) (anti-FLAG-tag Ab-immobilized beads) were used in this study. The beads used are described in each figure legend. When using S-protein agarose or Dynabeads Protein G, the washed beads were resuspended in SDS sample buffer and incubated at 95°C for 10 min. The precipitates were detected by WB. When using Anti DYKDDDDK tag Antibody Magnetic Beads, the washed beads were resuspended in SDS sample buffer without 2-mercaptoethanol and incubated overnight at 4°C. The beads were then removed, and 2-mercaptoethanol was added to the eluted sample followed by incubation at 95°C for 10 min. The samples were then subjected to WB.

### Electron microscopy and Southern blotting

Electron microscopy and Southern blotting were performed according to previously described methods (8).

### Statistics

The statistical significance was determined by one-way analysis of variance (ANOVA) followed by a Dunnett’s test and was evaluated using GraphPad Prism 7 software (GraphPad Software, CA, USA).

### Data availability

The underlying data and accession numbers are available in the main text. All other raw data are available upon request.

## Acknowledgements

The KSHV-BAC (BAC16) was kindly provided by Jae U. Jung (Cleveland Clinic Lerner Research Institute, USA). We thank Yoshihiko Fujioka (Osaka Medical and Pharmaceutical University, Japan) for electron microscopy analysis and Rimiko Okabe (Fujimuro Lab member) for assistance with data collection. Y.I. was supported by the Nagai Memorial Research Scholarship from the Pharmaceutical Society of Japan and the Japan Society for the Promotion of Science (JSPS) Research Fellowship for Young Scientists. This work was partially supported by grants from the SRF (2024G012) and the JSPS Grants-in-Aid for Scientific Research [18K06642 (M.F.) and JP22KJ2988 (Y.I.)].

---

**FIG S1.**
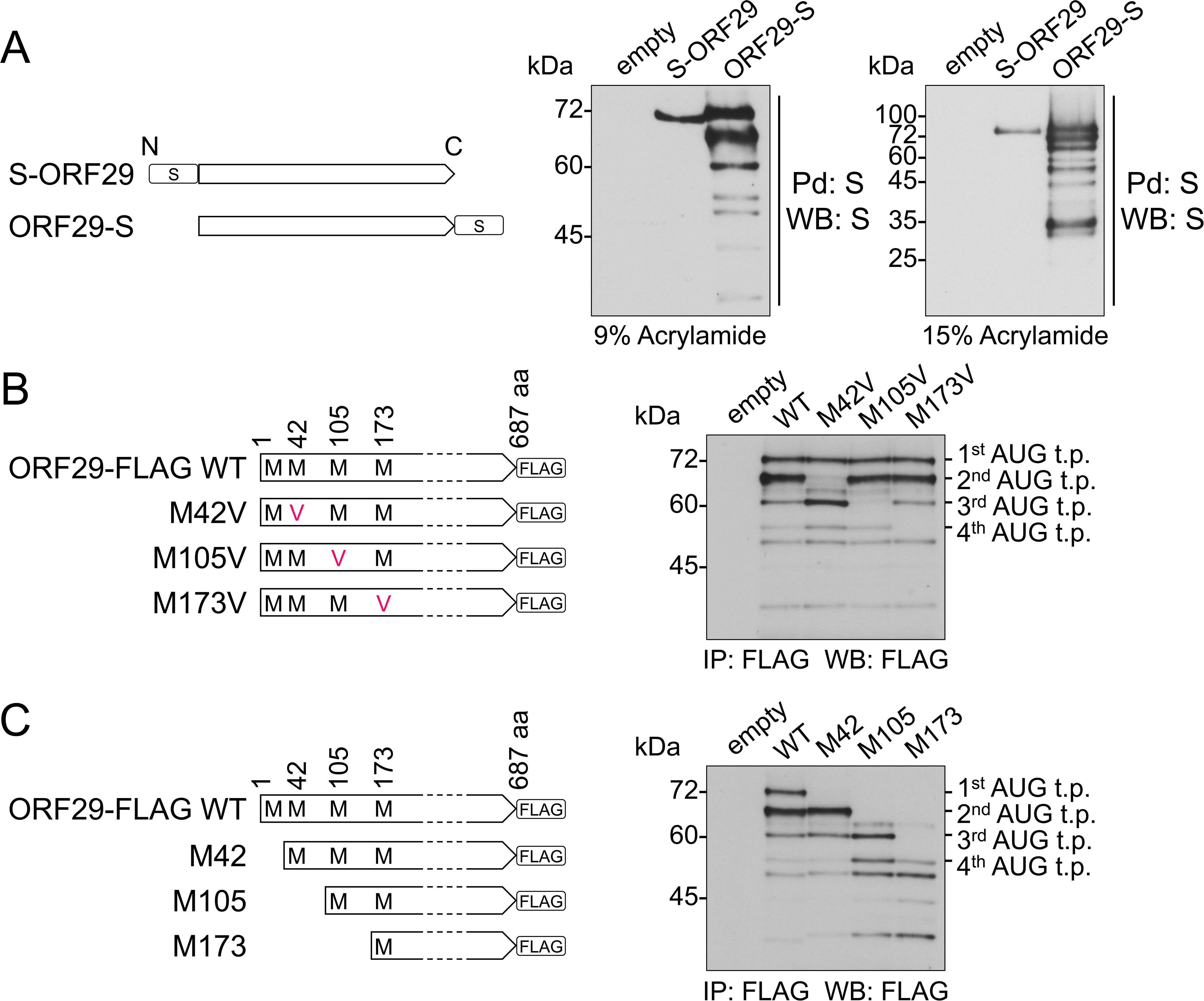
ORF29 short forms translated from the 2^nd^, 3^rd^, and 4^th^ AUG were expressed after transient transfection with the ORF29 plasmid. (A) N-terminal S-tagged ORF29 (S-ORF29) or C-terminal S-tagged ORF29 (ORF29-S) plasmid was transfected into HEK293T cells, and the cells were lysed in RIPA buffer. The lysates were subjected to Pd with S-protein agarose and were subjected to WB with anti-S Ab. (B, C) C-terminal FLAG-tagged WT ORF29 plasmid (ORF29-FLAG) or point-mutant ORF29 plasmid (ORF29-FLAG M42V, M105V, or M173V) or deletion-mutant ORF29 plasmid (ORF29-FLAG M42, M105, or M173) was transfected into HEK293T cells. The lysates were subjected to IP with anti-FLAG-tag Ab-immobilized beads and subjected to WB with anti-FLAG Ab. “t.p.” refers to translation product.

**FIG S2.**
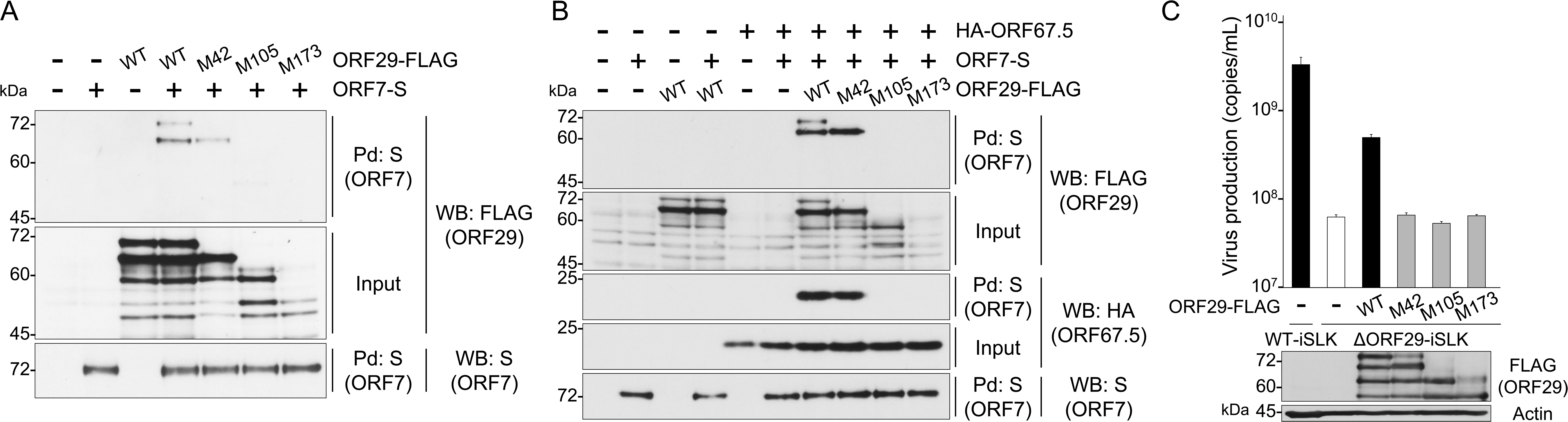
The ORF29 short forms do not contribute to ORF29 function. (A, B) HEK293T cells were transfected with the plasmids, and ORF7-S protein was Pd with S-protein agarose. The precipitated ORF7-S protein was analyzed by WB for detection of the ORF7-S-interacting molecule. (C) Each BAC-harboring iSLK cell line was transfected with the ORF29 short form plasmids. Simultaneously, the lytic phase was induced with Dox and SB for 72 h, and the culture supernatants were collected and treated with DNase. The number of encapsidated viral genome copies was quantified by qPCR. Bottom panel, expression of each exogenous ORF29 protein was confirmed by WB using an Ab directed against the FLAG-tag.

## References

1) Chang Y, Cesarman E, Pessin MS, Lee F, Culpepper J, Knowles DM, Moore PS. Identification of herpesvirus-like DNA sequences in AIDS-associated Kaposi’s sarcoma. Science. 1994 Dec 16;266(5192):1865–9. doi: 10.1126/science.7997879. PMID: 7997879.

2) Damania B, Cesarman E. 2022. Kaposi’s sarcoma herpesvirus, p 513–572. In Howley PM, Knipe DM (ed), Fields virology, 7th ed. Wolters Kluwer, Philadelphia, PA.

3) Russo JJ, Bohenzky RA, Chien MC, Chen J, Yan M, Maddalena D, Parry JP, Peruzzi D, Edelman IS, Chang Y, Moore PS. Nucleotide sequence of the Kaposi sarcoma-associated herpesvirus (HHV8). Proc Natl Acad Sci U S A. 1996 Dec 10;93(25):14862–7. doi: 10.1073/pnas.93.25.14862. PMID: 8962146; PMCID: PMC26227.

4) Heming JD, Conway JF, Homa FL. Herpesvirus Capsid Assembly and DNA Packaging. Adv Anat Embryol Cell Biol. 2017;223:119–142. doi: 10.1007/978-3-319-53168-7_6. PMID: 28528442; PMCID: PMC5548147.

5) Iwaisako Y, Fujimuro M. The Terminase Complex of Each Human Herpesvirus. Biol Pharm Bull. 2024;47(5):912–916. doi: 10.1248/bpb.b23-00717. PMID: 38692868.

6) Iwaisako Y, Watanabe T, Hanajiri M, Sekine Y, Fujimuro M. Kaposi’s Sarcoma-Associated Herpesvirus ORF7 Is Essential for Virus Production. Microorganisms. 2021 May 28;9(6):1169. doi: 10.3390/microorganisms9061169. PMID: 34071710; PMCID: PMC8228664.

7) Iwaisako Y, Watanabe T, Futo M, Okabe R, Sekine Y, Suzuki Y, Nakano T, Fujimuro M. The Contribution of Kaposi’s Sarcoma-Associated Herpesvirus ORF7 and Its Zinc-Finger Motif to Viral Genome Cleavage and Capsid Formation. J Virol. 2022 Sep 28;96(18):e0068422. doi: 10.1128/jvi.00684-22. Epub 2022 Sep 8. PMID: 36073924; PMCID: PMC9517700.

8) Iwaisako Y, Watanabe T, Suzuki Y, Nakano T, Fujimuro M. Kaposi’s Sarcoma-Associated Herpesvirus ORF67.5 Functions as a Component of the Terminase Complex. J Virol. 2023 Jun 29;97(6):e0047523. doi: 10.1128/jvi.00475-23. Epub 2023 Jun 5. PMID: 37272800; PMCID: PMC10308961.

9) Costa RH, Draper KG, Kelly TJ, Wagner EK. An unusual spliced herpes simplex virus type 1 transcript with sequence homology to Epstein-Barr virus DNA. J Virol. 1985 May;54(2):317–28. doi: 10.1128/JVI.54.2.317-328.1985. PMID: 2985801; PMCID: PMC254800.

10) Dolan A, Arbuckle M, McGeoch DJ. Sequence analysis of the splice junction in the transcript of herpes simplex virus type 1 gene UL15. Virus Res. 1991 Jun;20(1):97–104. doi: 10.1016/0168-1702(91)90064-3. PMID: 1656627.

11) Baines JD, Roizman B. The cDNA of UL15, a highly conserved herpes simplex virus 1 gene, effectively replaces the two exons of the wild-type virus. J Virol. 1992 Sep;66(9):5621–6. doi: 10.1128/JVI.66.9.5621-5626.1992. PMID: 1323715; PMCID: PMC289126.

12) Poon AP, Roizman B. Characterization of a temperature-sensitive mutant of the UL15 open reading frame of herpes simplex virus 1. J Virol. 1993 Aug;67(8):4497–503. doi: 10.1128/JVI.67.8.4497-4503.1993. PMID: 8331721; PMCID: PMC237833.

13) Baines JD, Poon AP, Rovnak J, Roizman B. The herpes simplex virus 1 UL15 gene encodes two proteins and is required for cleavage of genomic viral DNA. J Virol. 1994 Dec;68(12):8118–24. doi: 10.1128/JVI.68.12.8118-8124.1994. PMID: 7966602; PMCID: PMC237276.

14) Yu D, Sheaffer AK, Tenney DJ, Weller SK. Characterization of ICP6::lacZ insertion mutants of the UL15 gene of herpes simplex virus type 1 reveals the translation of two proteins. J Virol. 1997 Apr;71(4):2656–65. doi: 10.1128/JVI.71.4.2656-2665.1997. PMID: 9060618; PMCID: PMC191387.

15) Baines JD, Cunningham C, Nalwanga D, Davison A. The U(L)15 gene of herpes simplex virus type 1 contains within its second exon a novel open reading frame that is translated in frame with the U(L)15 gene product. J Virol. 1997 Apr;71(4):2666–73. doi: 10.1128/JVI.71.4.2666-2673.1997. PMID: 9060619; PMCID: PMC191388.

16) Yu D, Weller SK. Genetic analysis of the UL 15 gene locus for the putative terminase of herpes simplex virus type 1. Virology. 1998 Mar 30;243(1):32–44. doi: 10.1006/viro.1998.9041. PMID: 9527913.

17) Salmon B, Baines JD. Herpes simplex virus DNA cleavage and packaging: association of multiple forms of U(L)15-encoded proteins with B capsids requires at least the U(L)6, U(L)17, and U(L)28 genes. J Virol. 1998 Apr;72(4):3045–50. doi: 10.1128/JVI.72.4.3045-3050.1998. PMID: 9525627; PMCID: PMC109752.

18) Salmon B, Nalwanga D, Fan Y, Baines JD. Proteolytic cleavage of the amino terminus of the U(L)15 gene product of herpes simplex virus type 1 is coupled with maturation of viral DNA into unit-length genomes. J Virol. 1999 Oct;73(10):8338–48. doi: 10.1128/JVI.73.10.8338-8348.1999. PMID: 10482584; PMCID: PMC112851.

19) Renne R, Blackbourn D, Whitby D, Levy J, Ganem D. Limited transmission of Kaposi’s sarcoma-associated herpesvirus in cultured cells. J Virol. 1998 Jun;72(6):5182–8. doi: 10.1128/JVI.72.6.5182-5188.1998. PMID: 9573290; PMCID: PMC110093.

20) Arias C, Weisburd B, Stern-Ginossar N, Mercier A, Madrid AS, Bellare P, Holdorf M, Weissman JS, Ganem D. KSHV 2.0: a comprehensive annotation of the Kaposi’s sarcoma-associated herpesvirus genome using next-generation sequencing reveals novel genomic and functional features. PLoS Pathog. 2014 Jan;10(1):e1003847. doi: 10.1371/journal.ppat.1003847. Epub 2014 Jan 16. PMID: 24453964; PMCID: PMC3894221.

21) Miller JT, Zhao H, Masaoka T, Varnado B, Cornejo Castro EM, Marshall VA, Kouhestani K, Lynn AY, Aron KE, Xia A, Beutler JA, Hirsch DR, Tang L, Whitby D, Murelli RP, Le Grice SFJ. Sensitivity of the C-Terminal Nuclease Domain of Kaposi’s Sarcoma-Associated Herpesvirus ORF29 to Two Classes of Active-Site Ligands. Antimicrob Agents Chemother. 2018 Sep 24;62(10):e00233–18. doi: 10.1128/AAC.00233-18. PMID: 30061278; PMCID: PMC6153795.

22) McCollum CO, Didychuk AL, Liu D, Murray-Nerger LA, Cristea IM, Glaunsinger BA. The viral packaging motor potentiates Kaposi’s sarcoma-associated herpesvirus gene expression late in infection. PLoS Pathog. 2023 Apr 17;19(4):e1011163. doi: 10.1371/journal.ppat.1011163. PMID: 37068108; PMCID: PMC10138851.

23) Myoung J, Ganem D. Generation of a doxycycline-inducible KSHV producer cell line of endothelial origin: maintenance of tight latency with efficient reactivation upon induction. J Virol Methods. 2011 Jun;174(1-2):12–21. doi: 10.1016/j.jviromet.2011.03.012. Epub 2011 Mar 17. PMID: 21419799; PMCID: PMC3095772.

24) Brulois KF, Chang H, Lee AS, Ensser A, Wong LY, Toth Z, Lee SH, Lee HR, Myoung J, Ganem D, Oh TK, Kim JF, Gao SJ, Jung JU. Construction and manipulation of a new Kaposi’s sarcoma-associated herpesvirus bacterial artificial chromosome clone. J Virol. 2012 Sep;86(18):9708–20. doi: 10.1128/JVI.01019-12. Epub 2012 Jun 27. PMID: 22740391; PMCID: PMC3446615.

25) Deng B, O’Connor CM, Kedes DH, Zhou ZH. Cryo-electron tomography of Kaposi’s sarcoma-associated herpesvirus capsids reveals dynamic scaffolding structures essential to capsid assembly and maturation. J Struct Biol. 2008 Mar;161(3):419–27. doi: 10.1016/j.jsb.2007.10.016. Epub 2007 Nov 17. PMID: 18164626; PMCID: PMC2692512.

26) Gardner MR, Glaunsinger BA. Kaposi’s Sarcoma-Associated Herpesvirus ORF68 Is a DNA Binding Protein Required for Viral Genome Cleavage and Packaging. J Virol. 2018 Jul 31;92(16):e00840–18. doi: 10.1128/JVI.00840-18. PMID: 29875246; PMCID: PMC6069193.

27) Saveliev A, Zhu F, Yuan Y. Transcription mapping and expression patterns of genes in the major immediate-early region of Kaposi’s sarcoma-associated herpesvirus. Virology. 2002 Aug 1;299(2):301–14. doi: 10.1006/viro.2002.1561. PMID: 12202233.

28) Veronese BHS, Nguyen A, Patel K, Paulsen K, Ma Z. ORF48 is required for optimal lytic replication of Kaposi’s sarcoma-associated herpesvirus. PLoS Pathog. 2024 Aug 26;20(8):e1012081. doi: 10.1371/journal.ppat.1012081. PMID: 39186813; PMCID: PMC11379392.

29) Majerciak V, Alvarado-Hernandez B, Lobanov A, Cam M, Zheng ZM. Genome-wide regulation of KSHV RNA splicing by viral RNA-binding protein ORF57. PLoS Pathog. 2022 Jul 14;18(7):e1010311. doi: 10.1371/journal.ppat.1010311. PMID: 35834586; PMCID: PMC9321434.

30) AuCoin DP, Pari GS. The human herpesvirus-8 (Kaposi’s sarcoma-associated herpesvirus) ORF 40/41 region encodes two distinct transcripts. J Gen Virol. 2002 Jan;83(Pt 1):189–193. doi: 10.1099/0022-1317-83-1-189. PMID: 11752716.

31) AuCoin DP, Colletti KS, Cei SA, Papousková I, Tarrant M, Pari GS. Amplification of the Kaposi’s sarcoma-associated herpesvirus/human herpesvirus 8 lytic origin of DNA replication is dependent upon a cis-acting AT-rich region and an ORF50 response element and the trans-acting factors ORF50 (K-Rta) and K8 (K-bZIP). Virology. 2004 Jan 20;318(2):542–55. doi: 10.1016/j.virol.2003.10.016. PMID: 14972523.

32) Bruce AG, Barcy S, DiMaio T, Gan E, Garrigues HJ, Lagunoff M, Rose TM. Quantitative Analysis of the KSHV Transcriptome Following Primary Infection of Blood and Lymphatic Endothelial Cells. Pathogens. 2017 Mar 19;6(1):11. doi: 10.3390/pathogens6010011. PMID: 28335496; PMCID: PMC5371899.

33) Yang Y, Yang P, Wang N, Chen Z, Su D, Zhou ZH, Rao Z, Wang X. Architecture of the herpesvirus genome-packaging complex and implications for DNA translocation. Protein Cell. 2020 May;11(5):339–351. doi: 10.1007/s13238-020-00710-0. Epub 2020 Apr 23. PMID: 32328903; PMCID: PMC7196598.

34) Abbotts AP, Preston VG, Hughes M, Patel AH, Stow ND. Interaction of the herpes simplex virus type 1 packaging protein UL15 with full-length and deleted forms of the UL28 protein. J Gen Virol. 2000 Dec;81(Pt 12):2999–3009. doi: 10.1099/0022-1317-81-12-2999. PMID: 11086131.

35) Yang K, Baines JD. The putative terminase subunit of herpes simplex virus 1 encoded by UL28 is necessary and sufficient to mediate interaction between pUL15 and pUL33. J Virol. 2006 Jun;80(12):5733–9. doi: 10.1128/JVI.00125-06. PMID: 16731912; PMCID: PMC1472570.

36) Tischer BK, Smith GA, Osterrieder N. En passant mutagenesis: a two step markerless red recombination system. Methods Mol Biol. 2010;634:421–30. doi: 10.1007/978-1-60761-652-8_30. PMID: 20677001.

37) Yamaguchi T, Watanabe T, Iwaisako Y, Fujimuro M. Kaposi’s Sarcoma-Associated Herpesvirus ORF21 Enhances the Phosphorylation of MEK and the Infectivity of Progeny Virus. Int J Mol Sci. 2023 Jan 8;24(2):1238. doi: 10.3390/ijms24021238. PMID: 36674756; PMCID: PMC9867424.

